# Analysis of evolutionary dynamics and clonal architecture in prostate cancer

**DOI:** 10.1101/2023.03.23.533974

**Authors:** Jake R. Conway, Alok K. Tewari, Sabrina Y. Camp, Seunghun Han, Jett Crowdis, Meng Xiao He, Yaw A. Nyame, Saud H. AlDubayan, Nikolaus Schultz, Zoltan Szallasi, Mark M. Pomerantz, Matthew L. Freedman, Lawrence Fong, Peter S. Nelson, Myles Brown, Keyan Salari, Eliezer Van Allen

## Abstract

The extent to which clinical and genomic characteristics associate with prostate cancer clonal architecture, tumor evolution, and therapeutic response remains unclear. Here, we reconstructed the clonal architecture and evolutionary trajectories of 845 prostate cancer tumors with harmonized clinical and molecular data. We observed that tumors from patients who self-reported as Black had more linear and monoclonal architectures, despite these men having higher rates of biochemical recurrence. This finding contrasts with prior observations relating polyclonal architecture to adverse clinical outcomes. Additionally, we utilized a novel approach to mutational signature analysis that leverages clonal architecture to uncover additional cases of homologous recombination and mismatch repair deficiency in primary and metastatic tumors and link the origin of mutational signatures to specific subclones. Broadly, prostate cancer clonal architecture analysis reveals novel biological insights that may be immediately clinically actionable and provide multiple opportunities for subsequent investigation.

**Statement of significance:** Tumors from patients who self-reported as Black demonstrate linear and monoclonal evolutionary trajectories yet experience higher rates of biochemical recurrence. In addition, analysis of clonal and subclonal mutational signatures identifies additional tumors with potentially actionable alterations such as deficiencies in mismatch repair and homologous recombination.

## Introduction

Tumors consist of cell subpopulations that are characterized by a variety of features, including single nucleotide variants (SNVs), insertions and deletions (indels), copy number alterations (CNAs), structural variants (SVs), and epigenomic variation <u>(1)</u>. The aggregate of these subpopulations defines the clonal architecture and heterogeneity of a tumor. Whole-exome sequencing can provide a snapshot of these cell subpopulations at a point in time and space, and computational methods aimed at determining the number of tumor cell subpopulations and their relative cellular frequencies (“clonal architecture”), as well as inferring linear or branched phylogenetic evolutionary trajectories <u>(2–4)</u>, can improve our understanding of tumor evolution. Understanding the relationship between tumor clonal architecture and patient clinical characteristics may provide insights into disease trajectory and outcomes, as well as inform treatment decision-making <u>(5)</u>.

Prostate cancer (PC) is the second leading cause of cancer mortality in men <u>(6)</u>. Genomically-informed therapeutic options in advanced PC were limited until the approval of immune checkpoint blockade with pembrolizumab for tumors with deficient DNA mismatch repair (dMMR), microsatellite instability (MSI) <u>(7)</u>, or high tumor mutational burden, and PARP inhibition for prostate tumors harboring certain deleterious alterations in homologous recombination (HR) genes <u>(8,9)</u>. Despite their overall benefit, responses to these therapies are heterogeneous <u>(8–10)</u>, and little is known to date regarding how these clinical phenotypes and treatment responses relate to the clonal architecture of the tumors.

A prior study leveraged a large cohort of localized PC tumors (n = 293) to analyze associations between clonal architecture and clinical (e.g., Gleason score) or genomic (e.g., mutational signatures) covariates (11). In that cohort, polyclonality was associated with certain higher risk features such as PSA level and increased risk of biochemical recurrence after definitive therapy, and the mutational signature spectrum of a subset of primary tumors shifted over time from clock-like to that of homologous recombination deficiency (HRD). However, clonal architecture analyses to date have generally been limited to primary tumors, did not examine differences across self-identified race, and have not examined the clonality of mutational signatures. We hypothesized that clonal architecture analysis in a larger and more diverse cohort including non-White patients and metastatic tumors could improve our understanding of the genomic associations with clinical outcomes as well as clinical responses to emerging genomic treatment paradigms in metastatic PC. Herein, we paired harmonized molecular and clinical data from 845 primary and metastatic PC tumors <u>(12)</u> with novel computational methodologies to determine how clinical and genomic components relate to PC clonal architecture and evolutionary dynamics.

## Results

### Localized Prostate Cancer Clinical Risk Groups and Clonal Architecture

We first evaluated previously identified <u>(11)</u> associations between clonal architecture (Figure 1A) and evolutionary trajectories (Figure 1B) with clinical characteristics in localized primary PCs. All primary PCs in this cohort were treatment-naïve radical prostatectomy specimens. Based on phylogenetic reconstruction (Methods), tumors were defined as (1) monoclonal or polyclonal (the former if only a single cell cluster was identified), and (2) having evidence of linear or branched evolution (the former if each cell subpopulation had a maximum of 1 child subpopulation). Each cluster of mutations identified within a tumor is referred to as a cell subpopulation or clone. A “clonal” subpopulation refers to the truncal clone, defined by the group of mutations present in the greatest proportion of tumor cells. “Subclonal” thus refers to child clones arising from the truncal clone.

**Figure 1:**
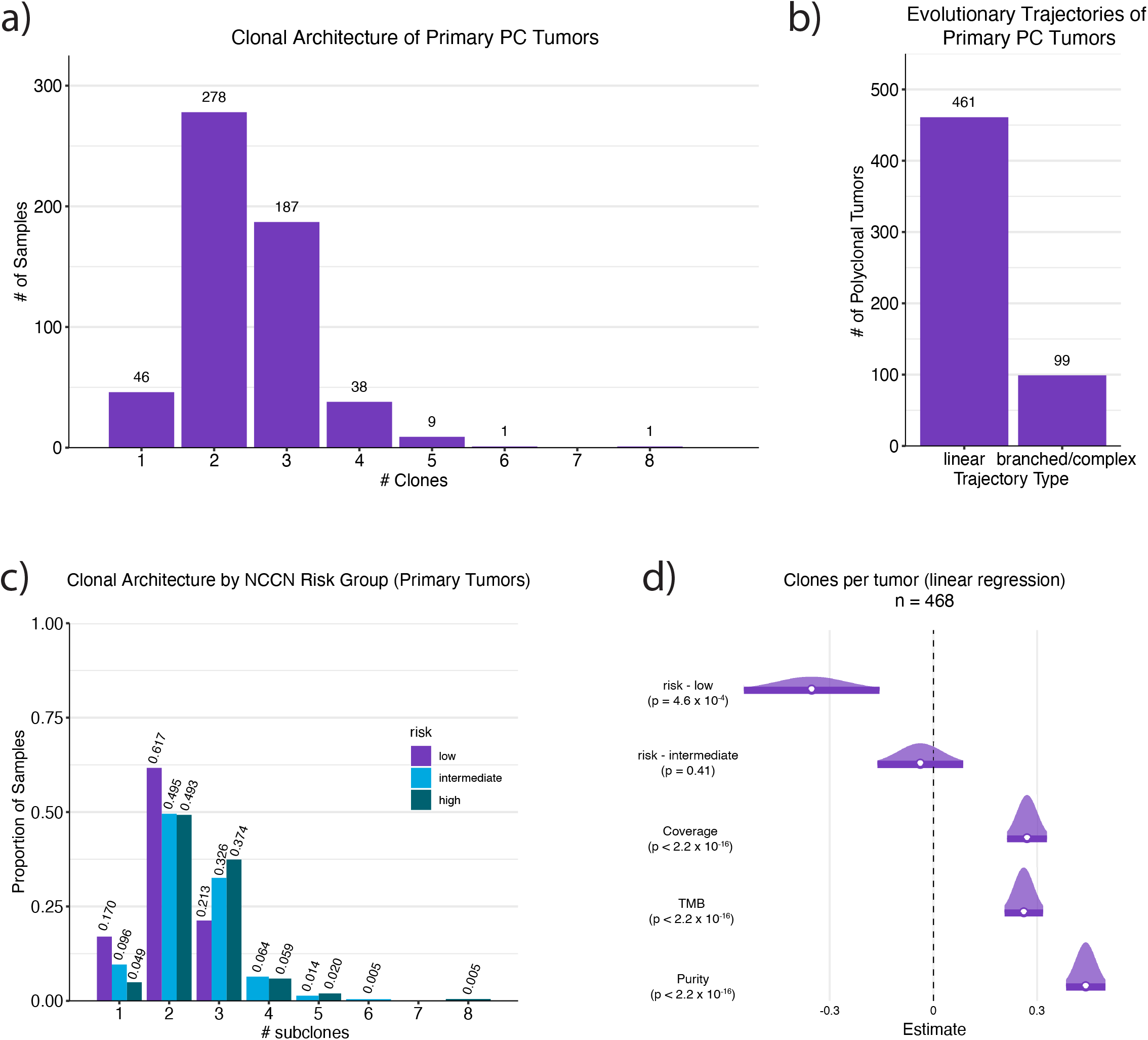
The clonal architecture and evolutionary trajectories of localized prostate cancer, and associations with clinical risk groups. **(a)** The distribution of the number of cell subpopulations (clonal architecture) per tumor across the cohort of 560 primary tumors. Of the 560 tumors, 46 (8%) were monoclonal, 278 (51%) were biclonal, and 236 (42%) were polyclonal. **(b)** Roughly three-quarters of PC tumors exhibited linear evolutionary trajectories rather than branched/complex evolutionary trajectories. **(c)** The distribution of the number of cell subpopulations (subclones) per tumor by risk group for all primary tumors with risk information in our cohort (n = 468). Low-risk tumors were significantly associated with being monoclonal relative to high-risk (Fisher’s; 95% CI = 1.26-11.88, OR = 3.93; p = 8.6 x 10^-3^), but not intermediate-risk (Fisher’s; 95% CI = 0.68-4.92, OR = 1.92; p = 0.19) tumors. **(d)** Low-risk tumors were also significantly associated with lower numbers of cell subpopulations per tumor after adjusting for confounding covariates, such as mutational burden, tumor purity, and coverage (linear regression, p = 4.6 x 10^-4^).

We did not detect a statistically significant association between clonal architecture and either age of PC onset (early: 55 years or younger; late: older than 55; univariate linear regression, p = 0.07) <u>(13)</u> or E26 transformation specific (ETS) fusion status in our cohort (n = 521; univariate linear regression, p = 0.1). However, we did identify significant associations between clonal architecture and National Comprehensive Cancer Network (NCCN) clinical risk categories (n = 468 primary tumors with risk information, Supplemental Tables 1-2). Approximately 17%, 10%, and 5% of low, intermediate, and high-risk primary tumors were classified as monoclonal, respectively. NCCN low-risk tumors were significantly associated with being monoclonal relative to high-risk tumors in univariate analysis (Figure 1C; Fisher’s exact test; 95% CI = 1.26-11.88, OR = 3.93; p = 8.6 x 10^-3^) and were significantly associated with fewer cell subpopulations even after adjusting for mutational burden, tumor purity, and coverage (Figure 1D; linear regression, p = 4.6 x 10^-4^). Relative to high-risk tumors, intermediate-risk tumors were not associated with monoclonal architecture (Fisher’s; 95% CI = 0.68-4.92, OR = 1.92; p = 0.19). Despite differences in the clonal architecture of tumors by risk group, there was no significant association between risk groups and evolutionary trajectories (linear vs. branched evolution; Fisher’s, p > 0.28, pairwise for all).

### Self-Reported Race and Clonal Architecture

The association between self-reported race and outcomes in men with prostate cancer is complex. Black men have substantially higher rates of prostate cancer incidence and mortality compared to other racial groups (14). These disparities are informed by social and structural determinants of equity and health, which impact a variety of socioeconomic, environmental, and health-related factors that influence prostate cancer biology and outcomes <u>(15,16)</u>. Due to the paucity of tumors from patients who self-report as Black in the majority of published genomic studies of PC, as well as limited information regarding the myriad of social factors that confound associations between race and outcome, the relationship between tumor genetics and inequities in clinical outcomes for Black patients is poorly understood <u>(17,18)</u>. To determine if the previously reported association between prostate cancer tumor polyclonality and clinical outcome extends to patients who self-report as Black, we performed clonal architecture analysis on the 560 primary PCs in our cohort, including 112 tumors from Black men (Supp. Table 1). We emphasize that our data set lacked any information on structural or socioeconomic factors, and therefore our analyses only included the following covariates: NCCN risk group (comprising stage, grade, and pre-treatment PSA) and sequencing characteristics (e.g., tumor coverage and purity).

Patients who self-reported as Black in our cohort were more likely to experience biochemical recurrence (Fisher’s, 26.9% vs. 18.7%, CI = 0.90-2.79; OR = 1.60, p = 0.08; Supp. Figure 1A), and this association was statistically significant after adjusting for clinical covariates (logistic regression, OR = 3.39, 95% CI = 1.78-6.56, p = 2.2 x 10^-4^). Patients who identified as Black had a significantly higher rate of early-onset PC development (Fishers, 36.6% vs. 27%, CI = 0.98 - 2.46, OR = 1.56, p = 0.048, Supp. Figure 1B), and had slightly lower median mutational burden (Mann-Whitney U, 1.04 mut/Mb vs. 1.27 mut/Mb, p = 1.13 x 10^-7^; Supp. Figure 1C). There was no difference in genomic instability, as measured by proportion genome altered, between tumors from Black and non-Black patients (Mann-Whitney U, 13.5% vs. 14.9%, p = 0.14), consistent with previous reports <u>(19)</u>.

We examined the association between self-reported race and various features of clonal architecture, adjusting our analyses using the available confounding covariates of clinical risk group, mutational burden, tumor purity, and sequencing coverage. Tumors from patients who self-reported as Black contained fewer clones (linear regression, p = 0.06, Figure 2A). Additionally, mutations in tumors from patients who self-reported as Black were found at higher cancer cell fractions (CCFs) (linear regression, p = 0.025, Figure 2B; Supp. Table 3). The overall CCFs of tumor cell subpopulations were also significantly higher in tumors from patients who self-reported as Black (Kolmogorov-Smirnov, p = 0.013, Figure 2C). The fraction of mutations classified as clonal per tumor was significantly higher in tumors from Black patients (Mann-Whitney U, p = 2.77 x 10^-6^, Figure 2D), even when removing monoclonal tumors from the analysis (Kolmogorov-Smirnov, p = 0.002), and this relationship remained significant when adjusting for the confounding covariates (linear regression, p = 0.034, Figure 2E; Supp. Table 3). When comparing tumor phylogenetics, tumors from patients who self-reported as Black were significantly associated with linear evolutionary trajectories (Fisher’s; 95% CI = 1.34-6.53; OR = 2.79; p = 0.003, Methods, after adjustment logistic regression, p = 0.036; Figure 2F).

**Figure 2:**
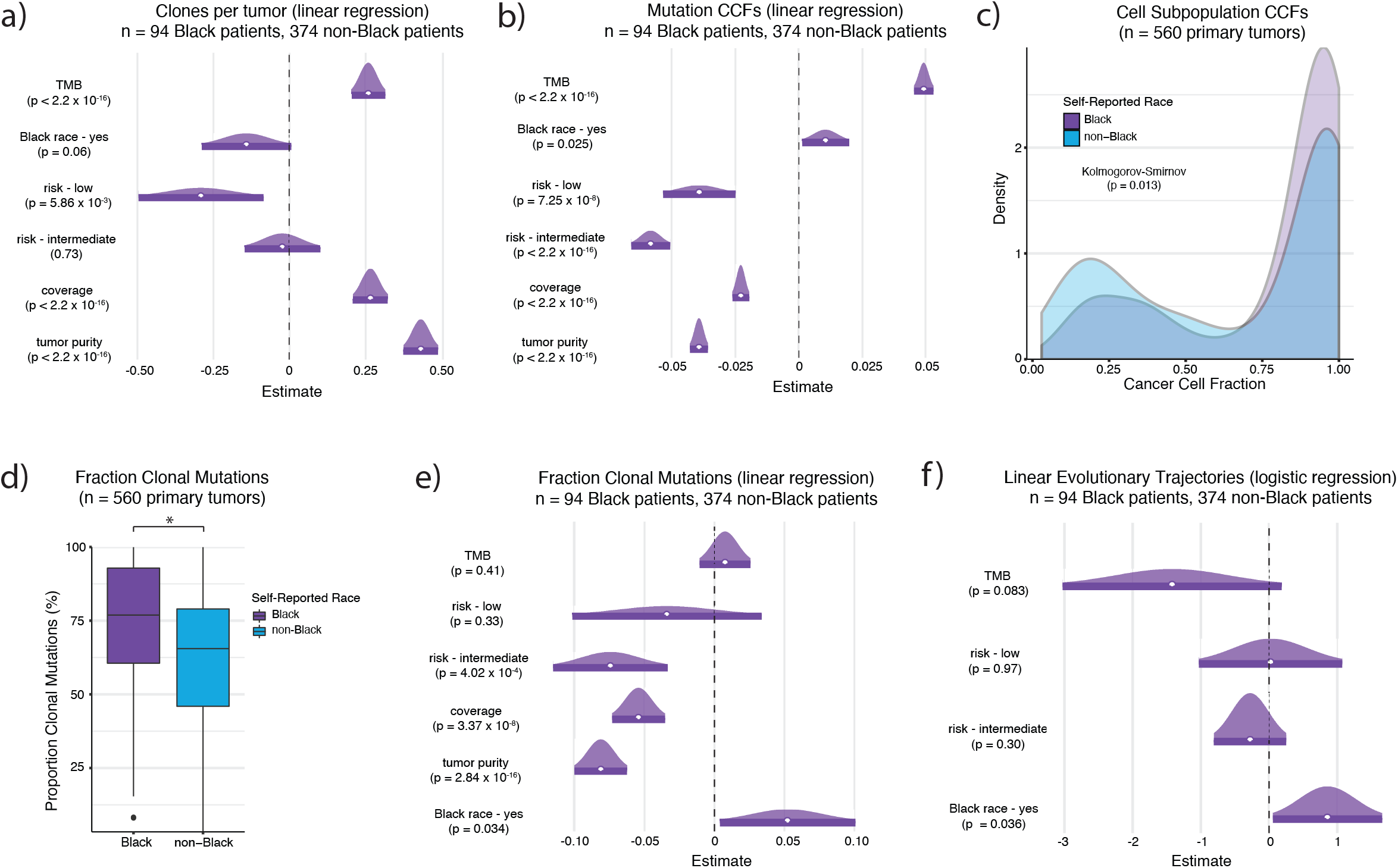
Primary prostate tumors from Black men exhibit more linear evolutionary trajectories. **(a)** Self-reported Black race was borderline significantly associated with fewer cell subpopulations (clones) per tumor (linear regression, p = 0.06), and **(b)** significantly associated with higher mutation cancer cell fractions (CCFs) compared to non-Black race (linear regression, p = 0.025) after correcting for confounding covariates. **(c)** The CCFs of cell subpopulations, not just individual mutations, was also significantly higher in tumors from Black patients compared to non-Black patients (Kolmogorov-Smirnov, p = 0.013). **(d)** The fraction of clonal mutations per tumor was significantly higher in samples from Black patients compared to samples from non-Black patients (Mann-Whitney U, p = 2.77 x 10^-6^), **(e)** even after correcting for confounding covariates (linear regression, p = 0.034). **(f)** Self-reported Black race was also associated with a higher frequency of linear evolutionary trajectories, rather than branched/complex evolutionary trajectories, compared to tumors from non-Black patients (logistic regression, p = 0.036).

To demonstrate that our findings were not a consequence of only sequencing coding regions, we performed the same analysis on a small cohort of primary PC whole-genomes from self-reported Black (n=7) and self-reported White (n=7) patients <u>(20)</u>. While mutation CCFs and the CCFs of tumor cell subpopulations were slightly higher in tumors from White patients, tumors from Black patients had fewer clones per tumor and a higher fraction of mutations classified as clonal per tumor (Supp. Figure 2A). In the context of this small cohort, these associations were not statistically significant after correcting for covariates (linear regression; fraction of clonal mutations: p = 0.08, Supp. Figure 2B; number of clones: p = 0.11). Although there were no monoclonal tumors in this cohort, 5 of 7 (71%) tumors from Black patients were bi-clonal compared to 3 of 7 (43%) tumors from White patients. Further, all tumors from Black patients (100%) had linear evolutionary trajectories compared to 3 of 7 (43%) tumors from White patients (Fisher’s; 95% CI = 0.85 - Inf; OR = Inf; p = 0.07; Figure 3).

**Figure 3:**
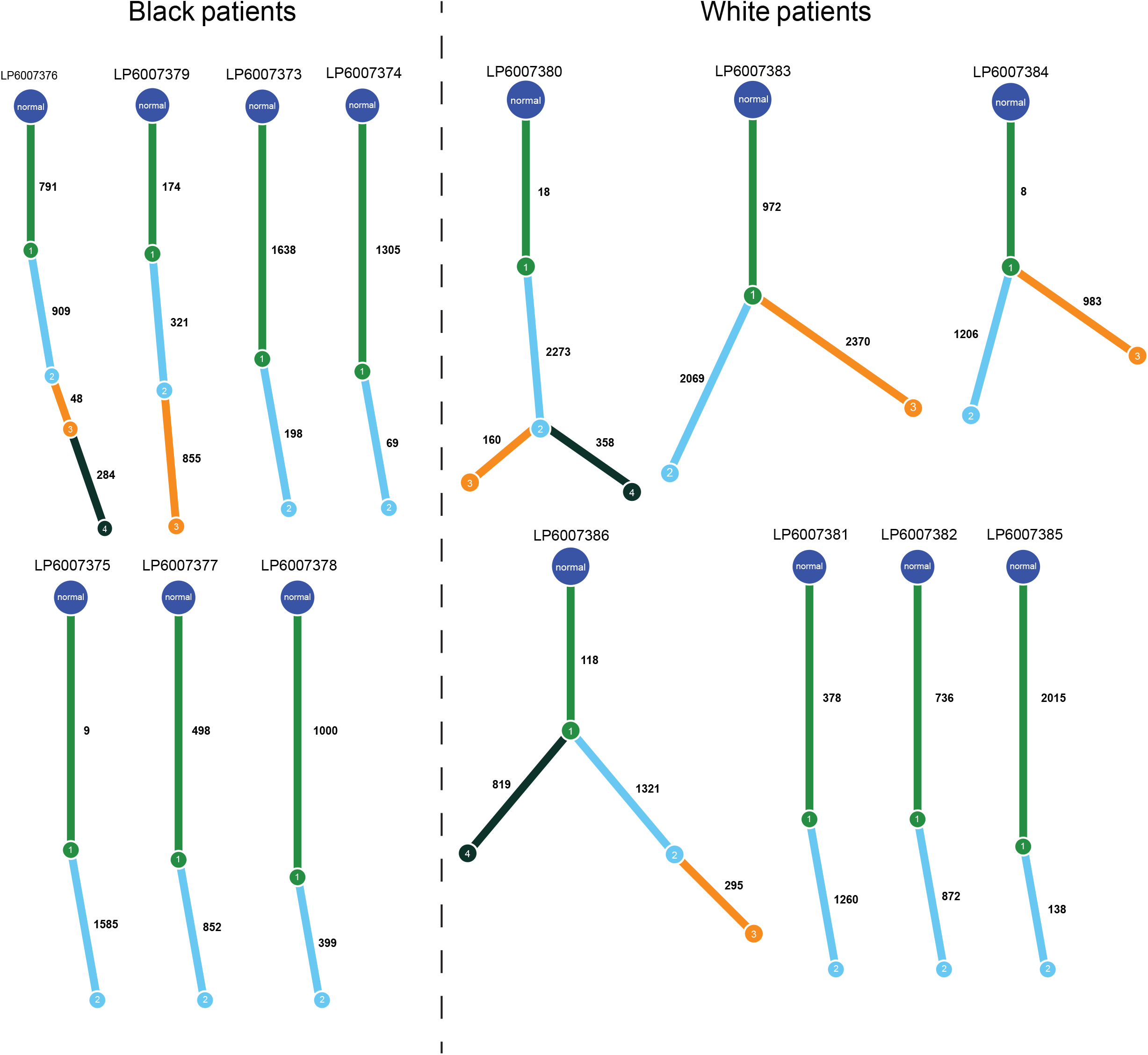
Evolutionary trajectories in whole genome sequencing samples from primary tumors in self-identified Black and White patients. A total of 2 out of 7 samples from Black patients had more than 2 cell subpopulations, compared to 4 out of 7 samples from White patients. In both cases from Black patients with more than 2 cell subpopulations, the tumor had a linear evolutionary trajectory, whereas all 4 cases from White patients with more than 2 cell subpopulations had branched evolutionary trajectories (Fisher’s, p = 0.07).

We also considered whether genetic ancestry inference could inform the relationship between self-reported race and clonal architecture. Consistent with a prior report (21), patient ancestry admixture did not add statistically independent information to the association between self-reported race and clonality in our cohort (p = 0.273 for interaction between self-reported race and AFR continental ancestry admixture). Recent work illustrates how continental-level genetic ancestry categories fail to capture the continuous genetic variation present in populations such as the United States, and that these categories are confounded by systems of racial classification (16,22). Therefore, we did not pursue ancestry analysis within the context of clonal architecture any further.

Together, these findings suggest an association between tumors from patients who self-reported as Black and linear trajectories as well as monoclonal genomic architecture. This finding contrasts with the positive correlation previously observed between polyclonality and more aggressive disease as measured by biochemical recurrence rate in a cohort of intermediate-risk PCs that did not include information about self-identified race <u>(11)</u>.

### Primary vs. Metastatic Disease and Clonal Architecture

We next aimed to compare clonal architecture and evolutionary trajectories between primary (N = 560) and metastatic (N = 285) PC tumors (Supp. Table 1). Primary tumors had significantly higher rates of monoclonality (Fisher’s; 10% vs. 2.5%; 95% CI = 1.93-11.40, OR = 4.32; p = 3.98 x 10^-5^) and fewer tumor cell subpopulations in general (Kolmogorov-Smirnov, p < 2.2 x 10^-16^; Supp. Figure 3B). When restricting to polyclonal tumors (> 2 subclones), to prevent the increased frequency of monoclonal and bi-clonal tumors in primary PCs from biasing the analysis, metastatic tumors were still significantly associated with branching patterns of evolution compared to primary tumors (Fisher’s; 68% vs. 52%; 95% CI = 1.22-3.05, OR = 1.92; p = 0.003), whereas primary tumors were significantly associated with linear evolution. When including monoclonal and bi-clonal tumors, over 80% of primary tumors displayed patterns of linear evolution, consistent with findings in whole-genomes (Supp. Figure 3C; Fisher’s; 82.4% vs. 67.4%; 95% CI = 1.6 – 3.18, OR = 2.25; p = 1.58 x 10^-6^) <u>(11)</u>.

We then evaluated clonal and subclonal significantly mutated genes (SMGs) in primary and metastatic PC tumors (q < 0.1). Six genes were identified exclusively as clonal SMGs in both primary and metastatic tumors (e.g., *APC*, *CDK12*, *ERF*, *FOXA1*, *SPOP*, *ZFHX3*), while 13 and 30 genes were identified exclusively as clonal SMGs in primary (e.g., *ATM*, *IDH1*, *SMARCA1*) or metastatic tumors (e.g., *AR*, *BRCA2*, *CUL3*), respectively. There was no overlap between subclonal primary and subclonal metastatic SMGs; however, the primary subclonal SMGs *PIK3CA*, *PTEN*, and *KDM6A* were identified as clonal SMGs in metastatic PCs (Supp. Figure 3D), supporting the hypothesis that alterations in the PI3K/AKT/mTOR pathway may increase the metastatic potential of primary PC tumors <u>(23–25)</u>. Indeed, the PI3K-AKT-mTOR signaling pathway was only identified via pathway overrepresentation analysis for primary subclonal SMGs and metastatic clonal SMGs (Methods; q < 0.05). Additional pathways overrepresented by clonal SMGs in metastatic samples included *AR* signaling, canonical *WNT* signaling, and the *RAC1*-*PAK1*-*p38*-*MMP2* pathway (q < 0.05). *AKT1 and NCOR1* were subclonal SMGs exclusively in primary tumors, and *AFF1* was a subclonal SMG exclusive to metastatic samples.

### Tumor-Level Mutational Signatures

Previous studies have shown that mutational signatures <u>(26,27)</u> appear and shift over the evolutionary trajectory of localized PC tumors <u>(11)</u>. Thus, we determined the mutational signatures present in each tumor and their association with clonal architecture. Using SigMA <u>(28)</u>, signature 3 was identified in 23.6% of metastatic PCs compared to 2% of primary PCs (Methods; Fisher’s; 95% CI = 7.79-32.59; OR = 15.21; p < 2.2 x 10^-16^, Figure 4A), and MSI associated signatures (signatures 6, 15, 20, and 26) were identified in 6.7% of metastatic PCs compared to 1.4% of primary PCs (Fisher’s; 95% CI = 2.01-13.10; OR = 4.89; p = 1 x 10^-4^), consistent with prior reports <u>(29–31)</u>. Conversely, clock-like signatures (signatures 1 and 5) were enriched as the dominant mutational process in primary PCs (Fisher’s; 96.6% vs. 70%; 95% CI = 7.14-21.84; OR = 12.21; p < 2.2 x 10^-16^). We orthogonally validated these classifications using scarHRD <u>(32)</u> to identify HRD-associated copy number events and MSIsensor <u>(33)</u> to detect somatic microsatellite changes (Supp. Figure 4A-C).

**Figure 4:**
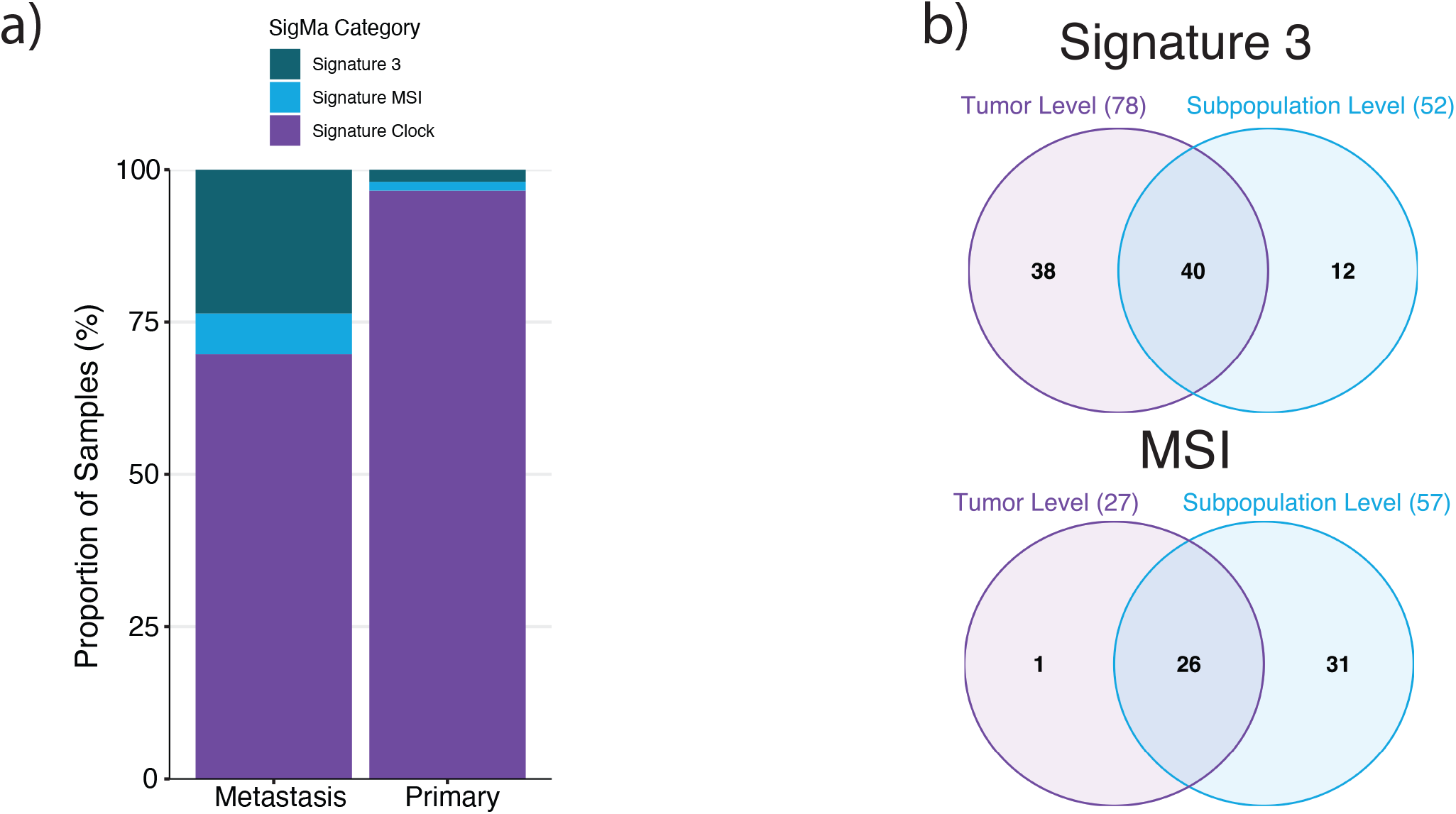
Mutational signature analysis at the cell subpopulation level can identify additional PC patients with HRD or MSI. **(a)** The proportion of metastatic and primary samples that have evidence of mutational signature 3 (associated with HRD) and MSI-associated mutational signatures. Activity of HRD and MSI-associated mutational signatures were more frequent in metastatic samples compared to primary samples. **(b)** The overlap between samples with evidence of mutational signature 3 and MSI-associated mutational signatures at the tumor and cell subpopulation levels. Out of 78 samples identified with signature 3 from running SigMA on all mutations in the tumor, 40 (51%) of those samples were identified as having signature 3 when running SigMA on the cell subpopulations. Conversely, 12 samples that were not classified as having signature 3 at the tumor level were identified as having signature 3 when running SigMA on the cell subpopulations. Similarly, while all but 1 sample identified with the MSI-associated signature at the tumor level were identified as having the MSI-associated signature at the cell subpopulation level, an 31 additional samples were identified as having MSI at the cell subpopulation level.

### Cell Subpopulation-Level Mutational Signatures

We next performed mutational signature analysis at the level of tumor cell subpopulations <u>(4)</u>. A total of 1439 clones across 829 tumors were powered for this analysis (Methods). Of the 78 tumors in our cohort with tumor level evidence of signature 3, 40 (51%) exhibited evidence of signature 3 at an individual clone level within the tumor, whereas 26/27 (96%) of samples with MSI at the aggregate tumor level showed evidence of MSI at the subpopulation level (Figure 4B). The discrepancy observed in the signature 3 tumors may be due in part to the reduction in power to call signature 3 when separating mutations into their respective clones. We utilized PyClone (2), a second phylogenetic reconstruction model, to determine the robustness of these signature calls and confirm their clonality (see Methods). There was over 93% agreement between the two methods for signature 3 and MSI calls both at the tumor and subclone level (Methods).

Of the 40 tumors with detectable signature 3 at both the overall tumor and cell subpopulation level, signature 3 was a clonal mutational process in 3 of 5 primary tumors (60%), and 31 of 35 metastatic tumors (89%). Of the 26 tumors with tumor and cell subpopulation level evidence of MSI, MSI was identified clonally in 6 of 8 primary tumors (75%), and 9 of 18 metastatic tumors (50%). Interestingly, 12 tumors categorized by clock-like mutational signatures at the tumor level demonstrated detectable signature 3 at the cell subpopulation level (Figure 4B), with the majority of these in the clonal cell subpopulation (83%; 1/2 primary, 9/10 metastatic). While tumors identified with signature 3 exclusively at the cell subpopulation level had higher numbers of HRD-associated copy number events than tumors identified with signature 3 exclusively at the whole tumor level, these differences were statistically non-significant (Supp. Figure 5). MSI was also detected in cell subpopulations of tumors overall categorized by clock-like mutational processes (n = 31, Figure 4B). In each of these tumors, MSI was detected at the subclonal level (1 primary, 30 metastatic). This high rate of subclonal MSI detection is consistent with results from the PCAWG cohort showing mismatch repair (MMR)-associated signatures preferentially result in subclonal mutations <u>(34)</u>. These results suggest that signature 3 and MSI classifications may be masked by more active or dominant mutational signatures, such as clock-like signatures in PC, when performing mutational signature analysis at the whole tumor level.

**Figure 5:**
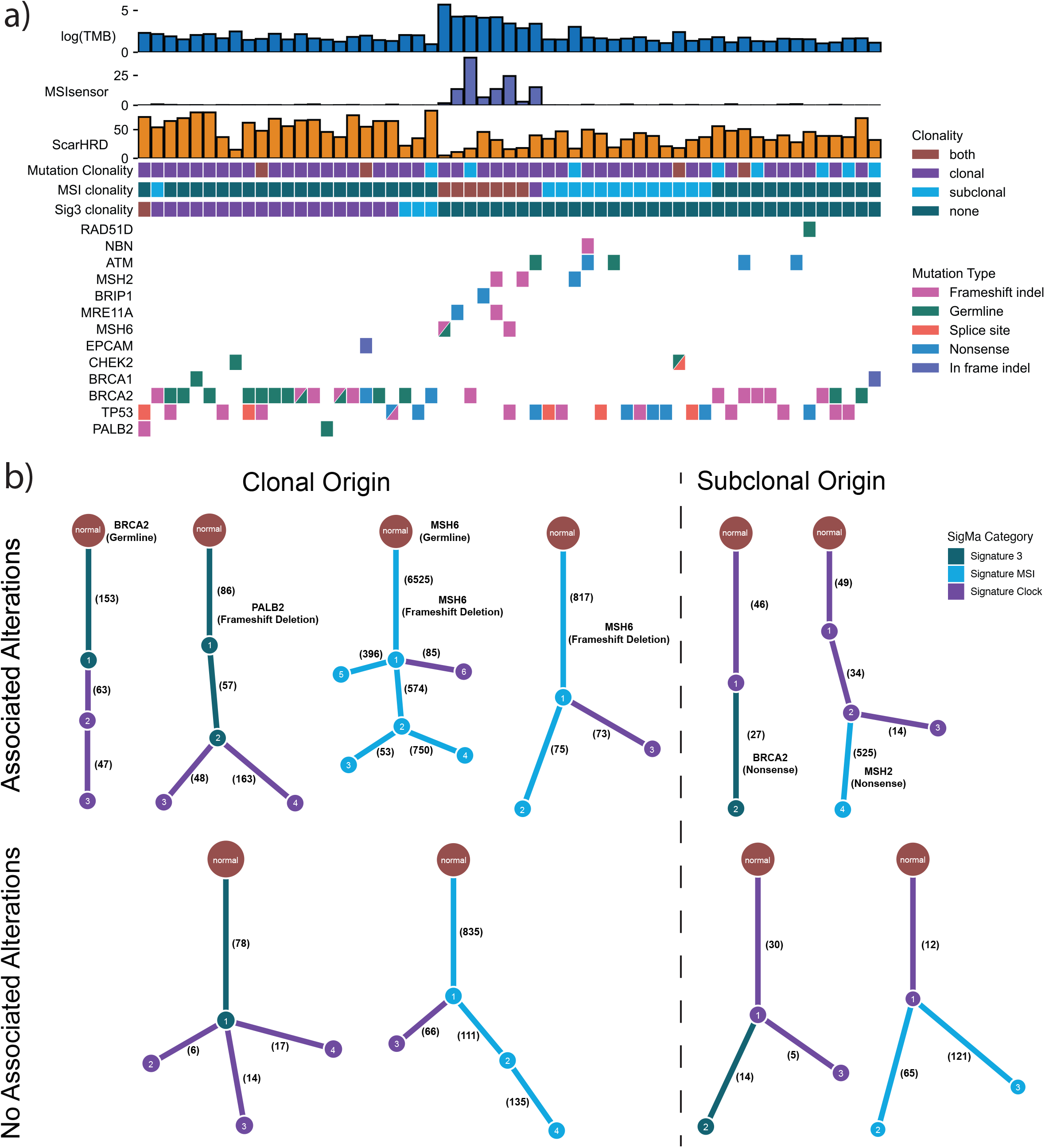
Linking the origin of mutational signatures to germline and somatic alterations and tracking how mutational signatures influence tumor evolution. **(a)** Comutation plot of putative LOF somatic and germline alterations in a curated set of DSB and MMR-associated genes. The comutation plot is annotated with the clonality of the alterations, clonality of signature 3 (associated with HRD), and clonality of MSI associated signatures. The comutation plot is also annotated with the number of HRD-associated CNA events (ScarHRD) per sample, as well as MSIsensor scores and mutational burden which are associated with MSI. **(b)** Phylogenetic trees from a subset of tumors included in the comutation plot showing how our novel approach can (**top**) link the origin of mutational signatures to somatic or germline alterations at the cell subpopulation level, and how (**top and bottom**) the mutational signatures influence the subclonal diversification over the tumors evolution. The subclone numbers indicate the inferred order in which those subclones occured. For example, subclone 2 is present at a higher CCF than subclone 3.

PC tumors with signature 3 present in the clonal subpopulation had significantly elevated numbers of LOH (loss of heterozygosity, Kolmogorov-Smirnov, p = 1.12 x 10^-10^; univariate logistic regression, p < 2.2 x 10^-16^), TAI (telomeric allelic imbalance, Mann-Whitney U, 23 vs. 4 events, p < 2.2 x 10^-16^), LST (large scale transitions, Mann-Whitney U, 23.5 vs. 8 events, p < 2.2 x 10^-16^), and total number of HRD-associated CNA (copy number alterations, Mann-Whitney U, 61 vs. 18 events, p < 2.2 x 10^-16^) events compared to PC tumors with no signature 3 (Supp. Figure 6A). However, PC tumors with clonal evidence of signature 3 had significantly elevated numbers of TAI (Mann-Whitney U, 23 vs.10 events, p = 8.4 x 10^-4^) events, but not LOH, LST, and total number of HRD-associated CNA events compared to PC tumors with subclonal signature 3 (Supp. Figure 6A). Tumors with subclonal evidence of signature 3 had significantly more LST (Mann-Whitney U, 17 vs. 8 events, p = 5.6 x 10^-3^) and total HRD-associated (Mann-Whitney U, 39 vs. 18 events, p = 0.01) CNA events compared to tumors with no signature 3, as well as borderline significant enrichment of LOH (Kolmogorov-Smirnov, p = 0.20; univariate logistic regression, p = 2.3 x 10^-9^) and TAI (Mann-Whitney U, 10 vs. 4 events, p = 0.058) events (Supp. Figure 6A).

Metastatic tumors with clonal evidence of MSI had significantly higher mutational burden and MSIsensor scores than metastatic tumors with subclonal MSI (TMB: Mann-Whitney U, 59.35 mut/Mb vs. 4.95 mut/Mb, p = 1.19 x 10^-9^; MSIsensor: Mann-Whitney U, 14.93 vs. 0.12, p = 3.63 x 10^-6^) or no MSI (TMB: Mann-Whitney U, 59.35 mut/Mb vs. 2.45 mut/Mb, p = 4.01 x 10^-7^; MSIsensor: Mann-Whitney U, 14.93 vs. 0.12, p = 2.83 x 10^-7^; Supp. Figure 6B-C). Of note, metastatic tumors with evidence of subclonal MSI had elevated mutational burden compared to metastatic tumors with no MSI signatures (Mann-Whitney U, 4.95 mut/Mb vs. 2.45 mut/Mb, p = 1.17 x 10^-9^, Supp. Figure 6C), but not higher MSIsensor scores (Mann-Whitney U, 0.12 vs. 0.12, p = 0.84; Supp. Figure 6B). Acknowledging the caveat that mutational signature detection depends on when a sample is procured within the evolutionary time of a tumor, these results suggest that the endogenous mutational process driving signature 3 may preferentially occur clonally, while the endogenous mutational process driving MSI may preferentially occur subclonally. Importantly, these results also suggest that subpopulation specific analyses can uncover additional samples with DNA repair defect-associated mutational signatures <u>(35,36)</u>.

### Mutational mechanisms of clonal and subclonal signature 3 and MSI in PC

We next aimed to leverage cell subpopulation level mutational signatures to identify genomic alterations associated with signature 3 and MSI (Figure 5A). Thus, we performed germline pathogenic mutation detection on a set of double-stranded break repair and MMR genes (Methods) and hypothesized that any germline alteration solely casual of signature 3 or MSI would manifest clonally. Of the 16 samples (all metastatic) with pathogenic germline *BRCA2* alterations, 10 (62.5%) showed clonal activity of signature 3 (Figure 5A-B, Supp. Figure 7A-B). Three of those 10 samples also had clonal somatic *BRCA2* mutations (1 missense, 2 frameshift deletions), and therefore signature 3 in these samples may be the result of biallelic loss. Conversely, none of the tumors with germline *ATM* alterations (0 of 10), and only 1 tumor with a *BRCA1* germline alteration (14%, 1/7) had clonal activity of signature 3. One of 2 samples with *PALB2* germline alterations, in this case a primary sample, also had clonal activity of signature 3. Additionally, 1 of 2 samples with *MSH6* germline alterations had clonal activity of MSI (Supp. Figure 7C-D), although this sample also had a clonal somatic *MSH6* mutation and therefore may be the result of a biallelic loss (Figure 5B).

We next identified cases where signature 3 and MSI may have been attributed to somatic alterations. Five of 15 tumors with only somatic putative loss of function (LOF) *BRCA2* mutations (i.e., no germline alterations) had signature 3. Three of these tumors had clonal *BRCA2* mutations, and 2 tumors had subclonal *BRCA2* mutations, with the onset of signature 3 being observed within the same cell subpopulation as the mutations (Figure 5B). Like the germline results, none of the tumors with putative LOF somatic *ATM* mutations had signature 3. One tumor with a clonal *BRCA1* mutation had clonal activity of signature 3, and 1 of 2 tumors with putative LOF somatic *PALB2* mutations (Figure 5B), each of which were clonal, had clonal evidence of signature 3.

Other than the MSI tumor with the *MSH6* biallelic loss, one other MSI tumor harbored a putative LOF *MSH6* somatic mutation (Figure 5B). The tumor with the *MSH6* double hit also had putative LOF somatic mutations in *MSH3* and *PMS1*, as well as missense mutations in 15 other MMR associated genes. Of note, this tumor had an intermediate MSIsensor score (1.81), despite having the highest mutational burden of any tumor in the cohort. All 3 samples with putative LOF somatic mutations in *MSH2* (2 clonal, 1 subclonal) had detectable MSI in the corresponding cell subpopulation (Figure 5A-B).

### Therapeutic implications of cell subpopulation mutational signature analysis

To determine the potential therapeutic implications of integrating cell subpopulation level mutational signatures with tumor level mutational signatures, we leveraged progression-free survival (PFS) data from patients in our cohort that were treated with PARP inhibitors (PARPi) (n=37 patients; Methods) <u>(37)</u>. Although not statistically significant in part due to the small sample size in this pilot cohort, univariate Cox PH analysis revealed that patients with tumor level (HR = 0.60, p = 0.31; Supp. Figure 8A) or cell subpopulation level (HR = 0.45, p = 0.095; Supp. Figure 8B) signature 3 had a lower hazard for progression when treated with PARPi, and this effect held when integrating both levels of signature calls (HR = 0.59, p = 0.20; Supp. Figure 8C). When correcting for prior treatment with radiation or systemic therapies, both cell subpopulation level signature 3 calls and the combination of tumor level and cell subpopulation level signature 3 calls were significantly associated with longer PFS (cell subpopulation level: HR = 0.34, p = 0.036; combined: HR = 0.39, p = 0.037; Supp. Figure 8D-F; Supp. Table 4), while tumor level signature 3 was borderline associated with improved PFS (HR = 0.33, p = 0.09). Thus, these results suggest that integrating cell subpopulation level mutational signatures with traditional approaches of identifying HRD patients can identify more patients that may benefit from PARPi.

To further validate the identification of cell subpopulation signature analysis and its therapeutic implications outside of the PC context, we applied this novel integrative approach to a cohort of ovarian cancer tumors treated with cisplatin chemotherapy <u>(38,39)</u> given the relevance of HRD alterations and associated therapeutics in this clinical context. We were able to recapitulate the association between signature 3 clonality and HRD-associated CNA events observed in PC (Supp. Figure 9A), and our approach identified additional patient tumors with signature 3 present compared to current approaches <u>(40,41)</u>. Furthermore, patients with signature 3 identified via our approach had improved PFS on cisplatin therapy (Supp. Figure 9B-E), further highlighting the potential expanded therapeutic impact of a clonal architecture-based approach to signature analysis for clinical use.

## Discussion

In this study, we revealed novel associations of genomic and clinical characteristics with clonal architecture through cancer cell fraction inference and phylogenetic reconstruction of 845 PC primary and metastatic tumor samples. Clinical covariates that were associated with the clonal architecture of PC tumors included clinical risk groups, self-reported race, and primary versus metastatic status. In contrast to the previously reported association between higher risk clinical parameters and more complex clonal architecture (11), we demonstrated that tumors from patients who self-reported as Black exhibit clonal architecture and evolutionary features associated with lower risk tumors. Our observations that primary tumors in Black patients are more monoclonal with linear genomic architectures yet have higher rates of biochemical recurrence, suggest at least two possibilities: (1) these tumors were screened for and detected at a time point after they had undergone rapid selective sweeps or punctuated evolution, and/or (2) other factors, potentially influenced entirely by structural, social and health effects, overshadow the potential positive prognostic value of linear clonal architecture seen in other patient populations.

Rapid selective sweeps in tumor evolution occur when all cell subpopulations are out-competed by one dominant clone <u>(5,42,43)</u>. The combination of mutations within this dominant clone may cause or collaborate with tumor epigenomic (44), transcriptomic, proteomic, and microenvironmental phenotypes that together contribute to higher rates of adverse clinical outcomes. The relative lower intratumoral heterogeneity in tumors from self-identified Black patients coupled with evidence of higher clonality genomic alteration events are akin to observations in clear cell renal cell carcinoma <u>(45)</u>, where these features were associated with worse outcomes compared to tumors with increased clonal diversity (i.e., more subclones) and lower clonality of alterations.

Together with prior publications suggesting that Black patients who undergo prostatectomy for PC have higher tumor volume relative to White patients (46), our data also reinforce the critical importance of more nuanced screening approaches and better healthcare resources and access for Black men. In the context of data illustrating equivalent if not better outcomes for Black patients with advanced PC when treated appropriately with standard of care therapies in relatively equal-access settings <u>(47–49),</u> more timely detection of PC in Black patients accompanied by prompt treatment in those with significant cancer and rigorous surveillance are critical to improving patient outcomes.

Our observations on self-reported race, genomic architecture, and outcome in localized PC have several additional caveats. Even in our larger cohort the number of Black patients is overall low, limiting the power to detect true population-level genomic differences. Our data was aggregated from patients who donated tissue samples primarily at certain academic medical centers through efforts such as The Cancer Genome Atlas, which do not fully represent the spectrum of PC in Black patients. Furthermore, our dataset lacks information on key associations with PC outcome such as health insurance access and zip code as well as other health factors such as obesity, tobacco exposure, and others (16,21). Analysis of more geographically and racially diverse patient populations and inclusion of information on known structural and social risk factors for prostate cancer incidence and mortality are vital for subsequent validation of our findings and other efforts to clarify the association between tumor biology and outcome. These efforts would require multiple strategies including engagement of Black patients, clinicians, and scientists to build biorepositories and cohorts that are enriched for geographically diverse populations of Black men.

Our novel approach integrating phylogenetic reconstruction <u>(4)</u> with mutational signature analysis <u>(28)</u> demonstrated that cell subpopulation level investigation can uncover both clonal and subclonal drivers as well as additional cases of HRD and MSI, which together may have therapeutic implications. Mutational significance analysis of clonal and subclonal mutations in primary and metastatic tumors revealed shared clonal drivers (e.g. *ERF*, *FOXA1* and *SPOP*) that may be important for tumor initiation and progression, as well as subclonal drivers in primary tumors that may confer metastatic potential and a selective advantage in metastatic sites that could be preferentially targeted (e.g. *PIK3CA*) <u>(23)</u>. We also find that associated genomic features, such as HRD-associated CNA events <u>(32)</u> and mutational burden <u>(50)</u>, show activity levels consistent with the clonality of the corresponding mutational signature. Specifically, there is a stepwise increase in the number of HRD-associated CNA events from tumors with no signature 3 to subclonal signature 3 to clonal signature 3, and a stepwise increase in mutational burden from tumors with no MSI signature to subclonal MSI signature to clonal MSI signature. Immunotherapy selection for PC patients based on MSIsensor scores <u>(51)</u> may miss subclonal MSI cases that can be captured through this analytical framework. For a subset of tumors, we further demonstrate the ability to link putative causal germline and somatic alterations to the origin of HRD or MSI <u>(29,30,52,53)</u> in the corresponding cell subpopulation. Additionally, the application of our novel integrative approach to PC tumors treated with PARPi and ovarian cancers treated with cisplatin chemotherapy identified more patients that appeared to benefit from these therapies, demonstrating the potential expanded therapeutic impact of our approach.

While the size of our cohort offers the statistical power necessary to find associations with clonal architecture and evolutionary trajectories, this study is limited by only having a single biopsy with exome sequencing per sample. Single-biopsy samples are more prone to sampling bias compared to multifocal biopsies, whereby mutation CCFs may deviate from their true value based on biopsy location (i.e., subclonal mutations presenting more clonally or subclonally) <u>(2,54,55)</u>. Though spatio-genomic data consisting of multi-focal biopsies from a single tumor are preferred for evolutionary analysis due to the increased power to resolve the true clonality of genomic alterations <u>(45,55)</u>, the number of PC patients with these data remains limited. To address this issue and increase the confidence of the timing (e.g., CCF or clonality status) of these events in PC, we leveraged the timing of each event across all samples and performed all analyses at the cohort level. Further, while PC whole-genome sequencing datasets do exist, they lack the breadth of clinical characteristics to fully interrogate self-reported race, HRD, and MSI but may augment these investigations prospectively. Future studies involving spatio-genomic profiling <u>(45,55)</u>, single-cell sequencing <u>(56–58)</u>, or long read sequencing <u>(59)</u> may further validate and inform the biology underlying these findings. Nevertheless, clonal architecture and evolutionary informed analysis of increasingly large cohorts of tumors will continue to reveal novel biological insights with immediate clinical potential, especially as clinical sequencing programs integrate such analyses into their workflows.

## Methods

### Cohort collection, quality control, and somatic variant calling

The somatic variants utilized in this study were taken from supplementary table 2 of Armenia *et. al* <u>(12)</u>. The cohort collection, quality control metrics, and somatic variant calling information can be found in the Methods of that study. However, in Armenia *et al* <u>(12)</u>, allelic copy number, purity, and ploidy information were determined using ABSOLUTE <u>(60)</u> in tumors where the purity was too low for FACETS <u>(61)</u> to output a solution. To remain consistent in this study, we re-ran FACETS and only kept tumors where FACETS produced a solution. Out of the original 1,013 PC tumors in the cohort, 845 produced a FACETS output.

### Clinical data

All clinical data was downloaded from the original published studies <u>(37,62–68)</u> Clinical data for the TCGA samples were taken from the original study <u>(63)</u>, and updated using the clinical data from the MC3 study <u>(39)</u> where applicable. Clinical risk groups were defined using the NCCN guidelines. Specifically, low risk: tumor stages T1-T2, grade group 1, and PSA < 10 ng/mL; intermediate risk: tumor stages T2b-T2c, grade group 2 or 3, or PSA 10-20 ng/mL, and no high risk features; high risk: tumor stage T3a or higher, grade group 4 or higher, or PSA > 20 ng/mL.

### Allelic copy number calling

Allelic CNAs were determined using FACETS<u>(61)</u>, which provides major and minor allele integer copy number values, tumor purity, tumor ploidy, and the cellular fraction of each copy number segment. The CCF of each CNA was calculated by adjusting the cellular fraction of the CNA by tumor purity.

### Calculation of mutation CCFs

Somatic mutation CCFs were calculated using the maximum likelihood method described in McGranahan *et al*.<u>(69)</u> using allelic copy number and tumor purity information from FACETS<u>(61)</u>. Here the maximum likelihood estimation of mutation CCFs are determined using a binomial distribution, taking into account tumor copy number, tumor purity, and variant allele frequency. This process is performed for the scenarios where the mutation occurs on either 1) the major allele, 2) the minor allele, or 3) a single allele copy, and the most likely CCF is chosen. For mutations that fall on normal ploidy segments, there is no difference in the CCF calculations for the mutation occurring on the major allele, minor allele, or a single copy of an allele. For mutations occurring on segments where both the major and minor allele copy numbers do not equal 1, the mutations with the highest likelihood of occurring on a single copy of an allele (rather than the major or minor allele) indicates that the mutation likely occurred after the CNA.

### Calculation of copy number alteration CCFs

The cellular frequency estimation column (“cf.em”) output by FACETS<u>(61)</u> is the fraction of all cells with a particular allelic copy number, including normal cells. To get the CCF of any non-diploid copy number segment the “cf.em” column was divided by the FACETS derived tumor purity.

### Phylogenetic reconstruction of tumor architecture

To reconstruct the clonal architecture of prostate cancer tumors we used the PhylogicNDT<u>(4)</u> Cluster module, which determines the number of tumor cell subpopulations and the respective assignment of each mutation to a cell subpopulation. The CCF annotated MAF file and FACETS-derived tumor purity were used as inputs to the clustering method. The outputs from the PhylogicNDT Cluster module were then used as inputs to the PhylogicNDT BuildTree module, which produces a series of phylogenetic trees ordered by likelihood. The phylogenetic trees with the highest likelihood were used in the analyses of this study, and were used to determine whether a tumor exhibited linear or branched evolution. Linear evolution is defined as phylogenetic reconstruction where each cell subpopulation in the tumor has a maximum of 1 child node. The number of clones per tumor were defined as the number of clusters identified by PhylogicNDT, and monoclonal tumors were defined as tumors where PhylogicNDT identified only a single cluster.

Although PyClone was not designed for use in whole-exome data <u>(2)</u> and tends to over cluster whole-exome mutation data <u>(70)</u>, it is one of the most highly used phylogenetic reconstruction methods. To validate the subclonal and clonal origin of mutational signatures (see Mutational signature analysis) we also ran PyClone with the following hyperparameters: burnin: 1000, density: pyclone_beta_binomial, init_method: connected, mesh_size: 101, num_iters: 10000, prior: major_copy_number, thin: 1. In cases where both methods were powered enough to call signatures at the cell subpopulation level, Pyclone identified MSI in all the same samples as PhylogicNDT (n=46; 100%), whereas Pyclone only identified signature 3 in 41 of 44 (93.2%) samples determined to have signature 3 via PhylogicNDT. Pyclone did not result in any additional MSI or signature 3 cases that weren’t detected by PhylogicNDT. Further, when both methods identified MSI and signature 3 in the same samples, MSI was identified at the same clonality (e.g. clonally or subclonally) in 43 of 46 (93.5%) samples, and signature 3 was identified at the same clonality in 40 of 41 (97.6%) samples.

### Mutational significance analysis

To perform mutation significance analysis we used MutSigCV2<u>(71)</u>, and classified SMGs as genes with an FDR corrected p-value < 0.1.

### Mutational signature analysis

Active mutational processes were determined using both the deconstructSigs<u>(40)</u> and SigMA <u>(28)</u> R packages. To run deconstructSigs we used the recommended, default parameters with the COSMIC (v2) signatures as the signatures reference. To run SigMA we set the data parameter equal to “seqcap” for whole-exome sequencing, the tumor_type parameter equal to “prost” for prostate adenocarcinoma, and check_msi parameter equal to TRUE to identify tumors with MMRd associated signatures. Default values were used for all other parameters. To run SigMA at the cell subpopulation level, we ran SigMA on the clusters of mutations output by PhylogicNDT<u>(4)</u>. In certain cases where there were too few mutations in a cluster (< 10), SigMA failed to produce an output for the cluster. Additionally, since running SigMA at the cell subpopulation level reduces the number of mutations input into SigMA, it may reduce the power to detect signature 3 and MSI in certain cases. Conversely, running at the cell subpopulation level enables the identification of signature 3 and MSI that may be confounded or overpowered by other mutational signatures at the tumor level.

While non-negative least squares (NNLS) methods (e.g. deconstructSigs)<u>(40)</u> are popular for identifying mutational signatures in individual samples, these methods are susceptible to increased rates of false positives in tumors with low mutational burden<u>(28)</u>, which is the case with many PC tumors. For instance, the observed prevalence of signature 3, associated with HRD, in primary PC whole-genomes was 5.8%, whereas deconstructSigs called signature 3 in 14.5% of primary PC tumors. A worse discrepancy was observed for mismatch repair deficient (MMRd) associated signatures. For this reason we reported mutational signature analysis using SigMA<u>(28)</u>, which utilizes non-negative matrix factorization (NMF), NNLS, and likelihood-based statistics combined with machine learning to classify known signatures across cancer types.

### Classification of clonal vs. subclonal mutations and mutational signatures

Mutations and mutational signatures were classified as clonal if they were identified in the truncal cluster of the tumor. Conversely, mutations and mutational signatures were classified as subclonal if they were not identified in the truncal cluster of the tumor. While mutations identified in the truncal cluster of the tumor are presumed to be present in every other cluster throughout the tumor, it is much more difficult to determine whether this is the case for mutational signatures. For instance, the identification of a mutational signature present in only the truncal cluster may still be active subclonally, but not identified due to power issues, the presence of a more active mutational signature, or that mutations generated by another signature were selected for. Conversely, mutations, CNAs or epigenetic alterations may cause a reversion of the mutational signature, specifically those caused by endogenous mutational processes such as HRD and MMRd.

### Calculation of HRD-associated CNA events (scores)

To calculate the number of LOH, TAI, and LST events in each tumor, we used the FACETS<u>(61)</u> allelic copy number calls as input to the scarHRD R package<u>(32)</u>. To determine the enrichment of these events in tumors classified with signature 3, we used the same statistical tests as the original papers the associations were discovered in. That is, Kolmogorov-Smirnov and univariate logistic regression for LOH events<u>(72)</u>, and Mann-Whitney U for both TAI and LST events<u>(73,74)</u>. Mann-Whitney U was also used to determine if there was a significant enrichment in the unweighted sum of events, and denote the significance values of all four scores (LOH, TAI, LST, unweighted sum) in the figures.

### Identification of mutations at microsatellites

To identify replication slippage variants at microsatellite regions and quantify the proportion that are somatic (also called MSIsensor score) we used MSIsensor<u>(33)</u>. PC tumors were characterized as having high, intermediate, and low MSIsensor scores using the same criteria as Abida *et. al* <u>(51)</u>.

### HRD and MMRd gene sets

The list of HRD and MMRd genes were curated from three prostate cancer specific studies: 1) Matteo *et al*. 2015<u>(29)</u>, 2) Pritchard *et al*. 2016<u>(30)</u>, and 3) de Bono *et al*. 2020<u>(52)</u>, as well as Polak *et al*. 2017<u>(53)</u>.

### Pathway overrepresentation analysis

We performed pathway overrepresentation analysis on genes identified as SMGs, and significantly amplified or deleted, using ConsensusPathDB (CPDB)<u>(75,76)</u>. We ran CPDB on December 3rd, 2019 with default parameters for pathway-based sets.

### Germline variant discovery

DeepVariant (v0.8.0)<u>(77,78)</u> was used to call SNVs and small deletions/duplications (indels) from whole-exome sequencing matched normal samples. Only high quality variants that were classified as “PASS” in the “FILTER” column were kept, and the CombineVariants module from GATK 3.7<u>(79)</u> was used to merge all of the high quality variants into a single Variant Call Format (VCF) file. The vt (v3.13)<u>(80)</u> tool was then used to decompose multiallelic variants, followed by normalization of variants. The high quality germline variants were annotated using VEP (v2)<u>(81)</u> with the publicly available GRCh37 cache file.

### Germline variant pathogenicity evaluation

High quality germline variants were evaluated for pathogenicity using publicly-available databases such as ClinVar and gene-specific databases, and classified according to the American College of Medical Genetics and Genomics and the Association of Molecular Pathology clinical-oriented guidelines<u>(82)</u>. Based on the evidence extracted from these resources, germline variants were classified into 5 categories: benign, likely benign, variants of unknown significance, likely pathogenic and pathogenic<u>(82)</u>. Additionally, truncating germline variants in genes that have yet been associated with a clinical phenotype, but are expected to disrupt the protein function, were classified as likely disruptive. Only germline variants classified as pathogenic, likely pathogenic, or likely disruptive were considered in the analysis.

### Ancestry inference

Hail (v0.2.39-ef87446bd1c7) was used to perform ancestry inference for each sample in our cohort. The “variant_qc” method was used on the combined cohort germline VCF to compute common variant statistics. This was followed by filtering out rare variants with an allele frequency less than 0.01, and variants that had a Hardy-Weinberg equilibrium p-value greater than 1 x 10^-6^. Additionally, we used the “ld_prune” method to filter out variants with a Spearman correlation threshold less than 0.1. The “hwe_normalized_pca” method was used to obtain the principal component analysis (PCA) eigenvalues and scores. To infer the ancestry of our samples, we also performed PCA on 1000 Genomes reference samples<u>(83,84)</u>, and trained a random forest classifier on the first 10 principal components to assign one of the five 1000 Genomes super populations (European, African, Admixed American, East Asian, and South Asian) to each of our samples.

### Population admixture estimation

Bcftools *mpileup* was used to generate a VCF of genotype likelihoods with the following options: 1) specifying not to skip anomalous read pairs (-A), 2) recalculating base alignment quality on the fly (-E), and 3) skipping indels (-I). We then called genotypes at local population ancestry reference sites from the 1000 Genomes Project (as opposed to super populations, see “Ancestry inference”; https://www.internationalgenome.org/category/population/) using Bcftools *call* with the option to call genotypes given alleles (-C), followed by removing duplicate genotype calls using Bcftools *norm*. The resulting VCF was used to as input to PLINK (v1.9)<u>(85)</u> *--make-bed*, which was subsequently used as input to fastNGSadmix<u>(86)</u> to determine the admixture proportions for the 1) Utah Residents (CEPH) with Northern and Western European Ancestry (CEU), 2) Han Chinese in Beijing, China (CHB), 3) Yoruba in Ibadan, Nigeria (YRI), and 4) Peruvians from Lima, Peru (PEL) populations.

### Survival analysis

To demonstrate the improved clinical utility of using SigMA<u>(28)</u>, and determine the therapeutic implication of cell subpopulation level mutational signatures, we performed survival analysis on a subset of patients in our cohort that received PARPi (Olaparib or Veliparib)<u>(37)</u>. In the multivariate model we corrected for whether or not the patient also received prior radiation, hormone therapy, chemotherapy, and immune-related therapy (i.e. yes or no). To validate the presence of signature 3 at both the tumor and cell subpopulation levels, and determine if the associations apply in other cancer contexts, we leveraged data from the TCGA ovarian cancer cohort<u>(38,39)</u>, and restricted our analysis to ovarian tumors treated with platinum-based chemotherapy. The drug information, number of platinum-based chemotherapy cycles, patient age, and tumor stage information were downloaded from FireCloud (https://portal.firecloud.org/#workspaces/broad-firecloud-tcga/TCGA_OV_ControlledAccess_V1-0_DATA). The ovarian cancer mutation calls and survival data were downloaded from the MC3 study<u>(39)</u>. To determine if there are significant differences between the survival curves of two or more groups we used the log-rank test from the survival R package. To evaluate whether any covariates were confounding the associations identified in the Kaplan-Meier analyses, we also performed Cox proportional hazards analysis (using the survival R package) correcting for these covariates.

### Clonal architecture and evolutionary dynamics in WGS cohort

Somatic mutation VCFs for African ancestry (n=7) and European (n=7) ancestry WGS samples from Petrovics *et al*<u>(20)</u> were obtained directly from the authors, and we restricted the somatic mutation call set to those classified as high-confidence in the original study. Mutation CCFs were calculated as described in “Calculation of mutation CCFs’’ using FACETS-derived allelic copy number calls and the histologically defined tumor purities from the original study. The clonal architecture and evolutionary trajectories of these tumors were determined using PhylogicNDT<u>(4)</u> as described in “Phylogenetic reconstruction of tumor architecture”.

## Acknowledgements

We gratefully acknowledge insightful commentary from Alham Sadaat and René Salazar at the Broad Institute. This work was supported by the Prostate Cancer Foundation (Young Investigator Awards to A.Tewari, S. AlDubayan and K. Salari, PCF-Movember Challenge Award to E. Van Allen and P. Nelson) and Urology Care Foundation Research Scholar Award (K. Salari). Support was also provided by the Mark Foundation Emerging Leader Award (E. Van Allen), the Department of Defense Prostate Cancer Research Program (W81XWH-20-1-0032 to A. Tewari, W81XWH-21-1-0084 and W81XWH-22-1-0455 to S. AlDubayan, W81XWH-21-1-0264 to P.Nelson and W81XWH-21-PCRP-DSA to E. Van Allen), and the National Institute of Health (U01 CA233100, R01 CA227388, P50CA097186, and P01 CA228696).

## Table Legends

**Supplementary Table 1:** Clinical data for 845 samples

**Supplementary Table 2:** PhylogicNDT results for primary PC tumors

**Supplementary Table 3:** Univariate and multivariate analysis results for associations with clonal architecture features

**Supplementary Table 4:** Univariate and multivariate COX PH models for PARPi treated PC patients

## Figure Legends

**Supplementary Figure 1:**
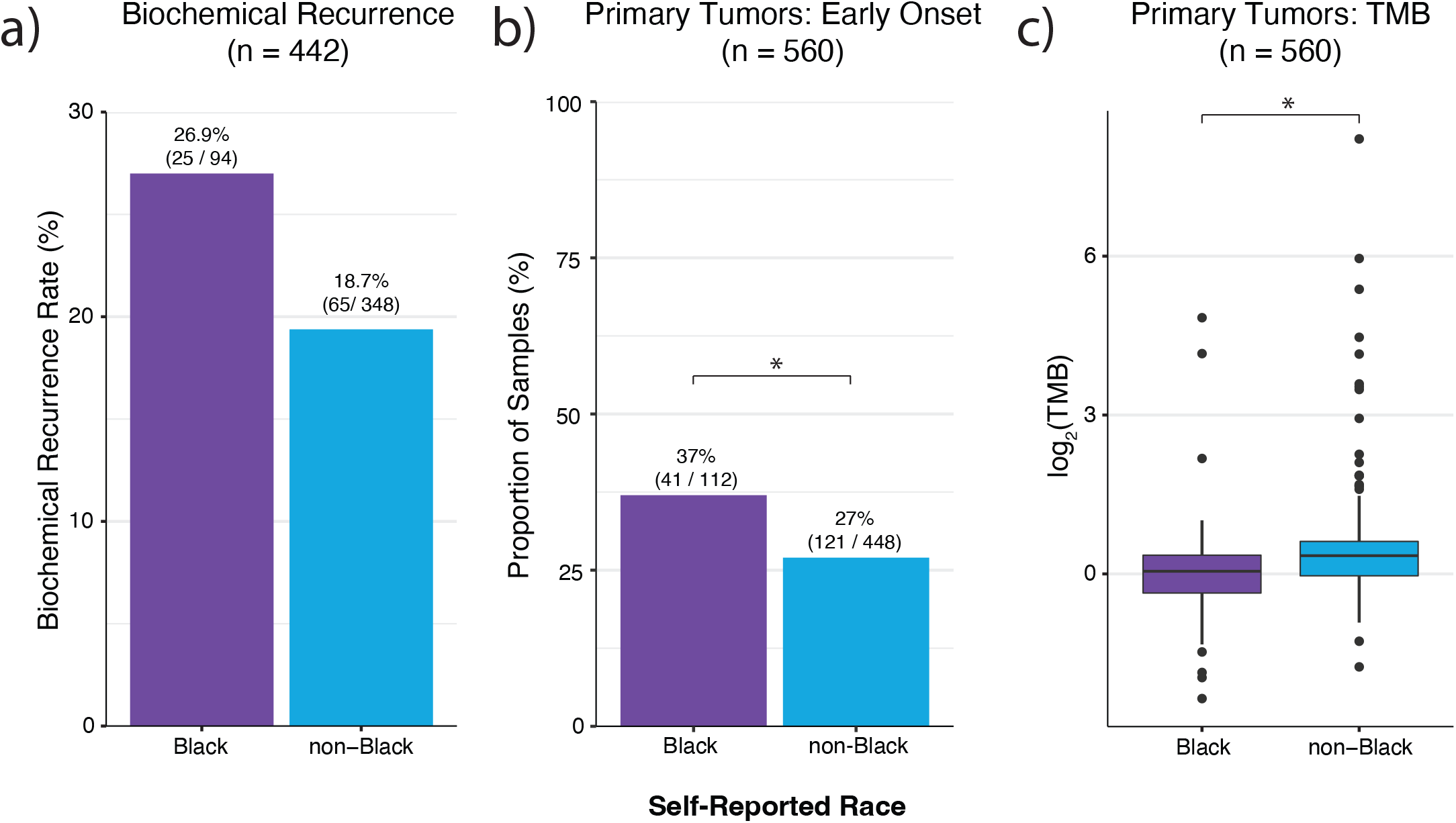
Clinical and Genomic Characteristics associated with self-reported race in PC. **(a)** The rate of biochemical recurrence between tumors from Black and non-Black patients with information on biochemical recurrence status (n=442). Consistent with previous studies, the rate of biochemical recurrence was higher in Black compared to non-Black patients (Fisher’s exact test, p = 0.08, logistic regression adjusting for clinical covariates, p = 2.2 x 10^-4^). **(b)** The proportion of early onset (<= 55 years old) tumors in our cohort between Black and non-Black patient tumors. Consistent with previous studies, Black patient samples were associated with a higher rate of early onset tumors (Fisher’s exact test, p = 0.048). **(c)** The distribution of TMB between Black and non-Black localized tumor samples. Non-Black patient samples were associated with a slightly higher TMB in our cohort (Mann-Whitney U, p = 1.13 x 10^-7^).

**Supplementary Figure 2:**
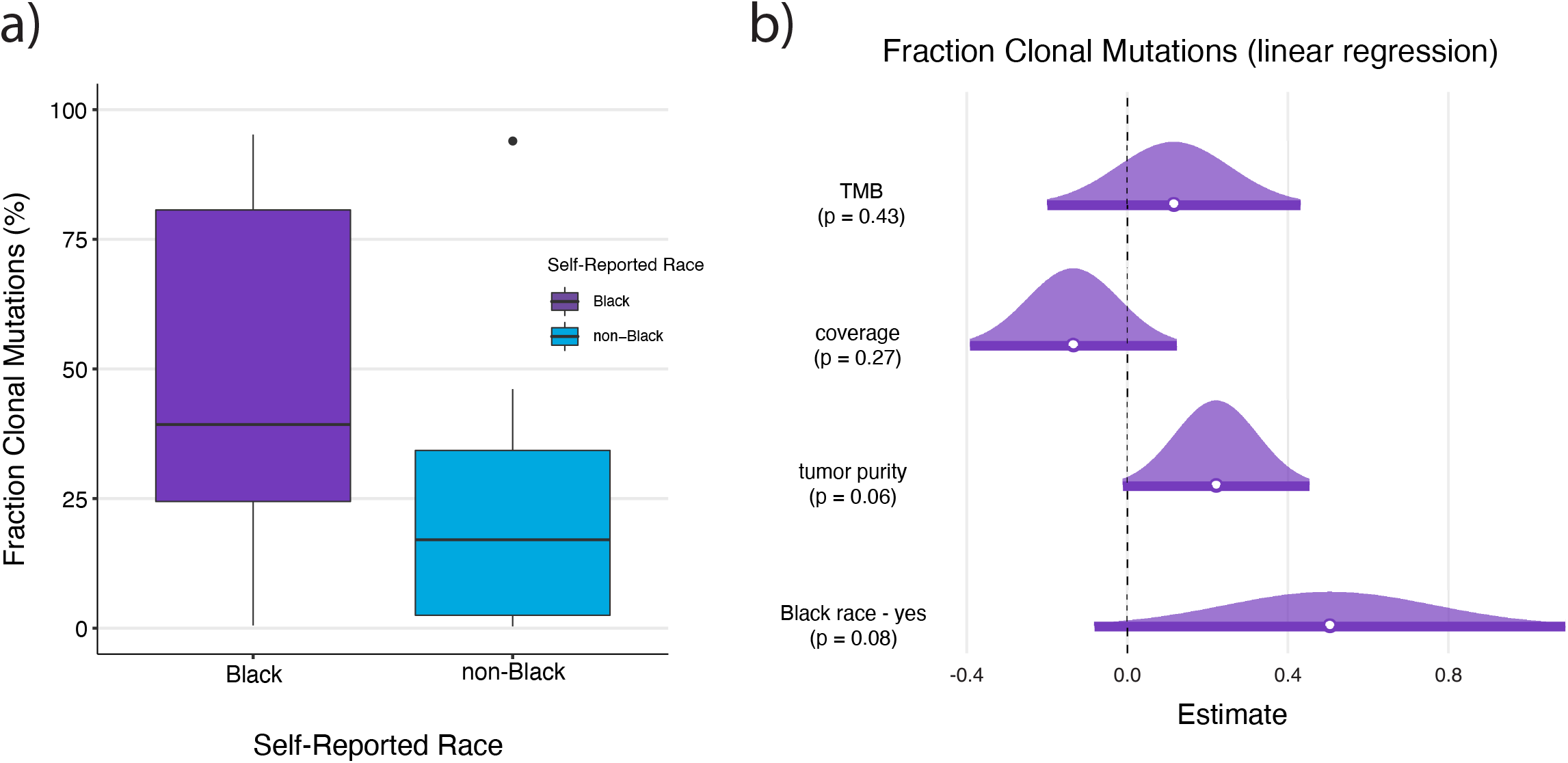
Whole genome sequencing validation of preferential punctuated evolutionary trajectories in tumors from Black patients. **(a)** The distribution of the fraction of clonal mutations in each whole genome sequenced tumor based on self-reported race (n = 7 Black patients, n = 7 White patients). Although non-significant (Mann-Whitney U, 39.3% vs. 17.1%, p = 0.32), Black patient tumors exhibited higher fractions of mutations in the clonal cluster. **(b)** After correcting for confounding covariates such as tumor mutational burden, sequencing coverage, and tumor purity, tumors from self-reported Black patients had higher proportion of mutations in the clonal cluster (multivariate linear regression, p = 0.08).

**Supplementary Figure 3:**
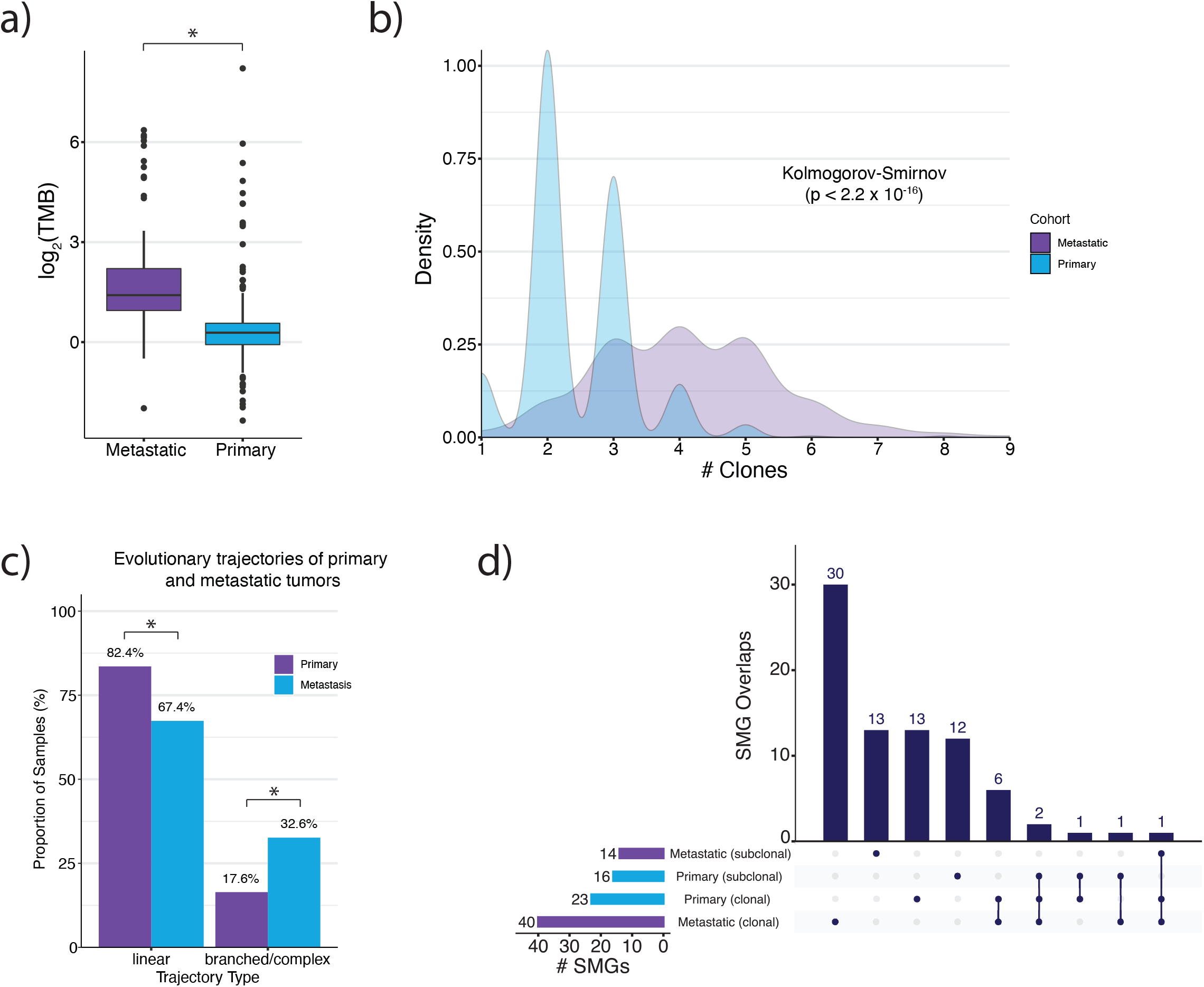
Primary and metastatic PC tumors are associated with distinct genomic and clonal architecture properties. **(a)** The distribution of tumor mutational burden (TMB) between primary and metastatic samples in our cohort (n = 845). The median TMB observed in the metastatic samples was more than two-fold higher than the TMB observed in primary samples (Mann-Whitney U, 2.66 mut/Mb vs. 1.22 mut/Mb, p < 2.2 x 10^-16^). **(b)** The distribution of the number of cell subpopulations (clones) between primary and metastatic samples. Metastatic samples were associated with having significantly more cell subpopulations than primary samples (Kolmogorov-Smirnov, p p < 2.2 x 10^-16^). Similarly, the rate of monoclonality in metastatic samples was 10% compared to 2.5% in primary samples (Fisher’s, p = 3.98 x 10^-5^). **(c)** Primary tumors are associated with a higher rate of linear evolutionary trajectories compared to metastatic tumors (Fisher’s; 95% CI = 1.6 - 3.18, OR = 2.25; p = 1.58 x 10^-6^). **(d)** The overlap between clonal and subclonal primary and metastatic driver genes identified via mutational significance analysis with MutSigCV2.

**Supplementary Figure 4:**
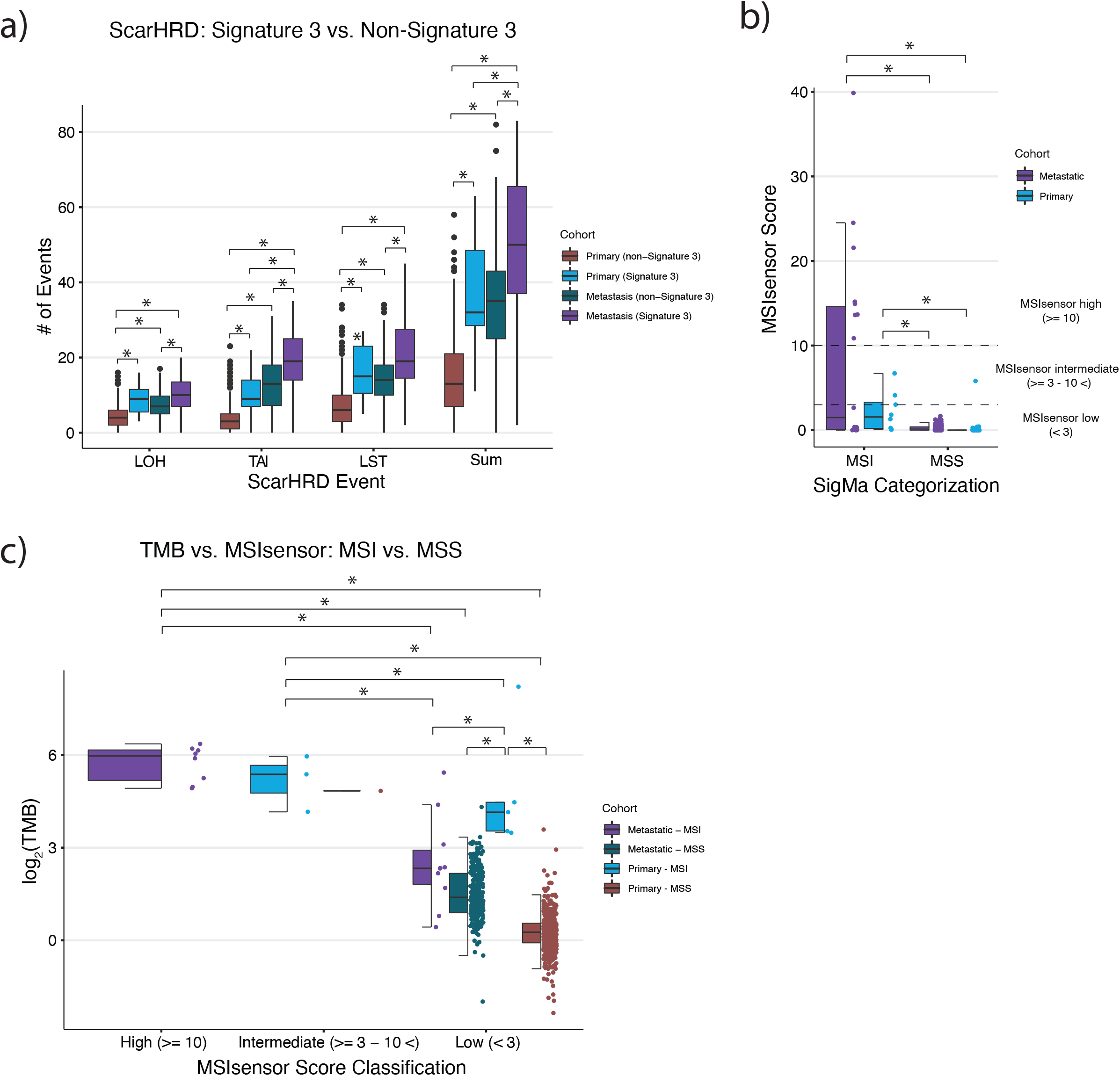
Mutational signatures and their associations with genomic events in PC tumors. **(a)** The distribution of homologous recombination deficiency (HRD)-associated copy number events between primary and metastatic samples with and without evidence of mutational signature 3. Metastatic samples with signature 3 were associated with higher numbers of HRD-associated copy number events compared to metastatic samples without signature 3, and primary samples with signature 3 were associated with higher numbers of HRD-associated copy number events compared to primary samples without signature 3. Additionally, metastatic samples with signature 3 had higher numbers of HRD-associated copy number events compared to primary samples with signature 3. **(b)** The distribution of MSIsensor scores between primary and metastatic samples with and without MSI-associated mutational signatures. Metastatic samples with MSI-associated mutational signatures had higher MSIsensor scores than metastatic samples without MSI-associated mutational signatures, and primary samples with MSI-associated mutational signatures had higher MSIsensor scores than primary samples without MSI-associated signatures. Additionally, metastatic samples with MSI-associated mutational signatures had higher MSIsensor scores compared to primary tumors with MSI-associated mutational signatures. **(c)** The distribution of mutational burden between primary and metastatic samples with and without MSI-associated mutational signatures. Asterisks denote statistical significance via Mann-Whitney U tests.

**Supplementary Figure 5:**
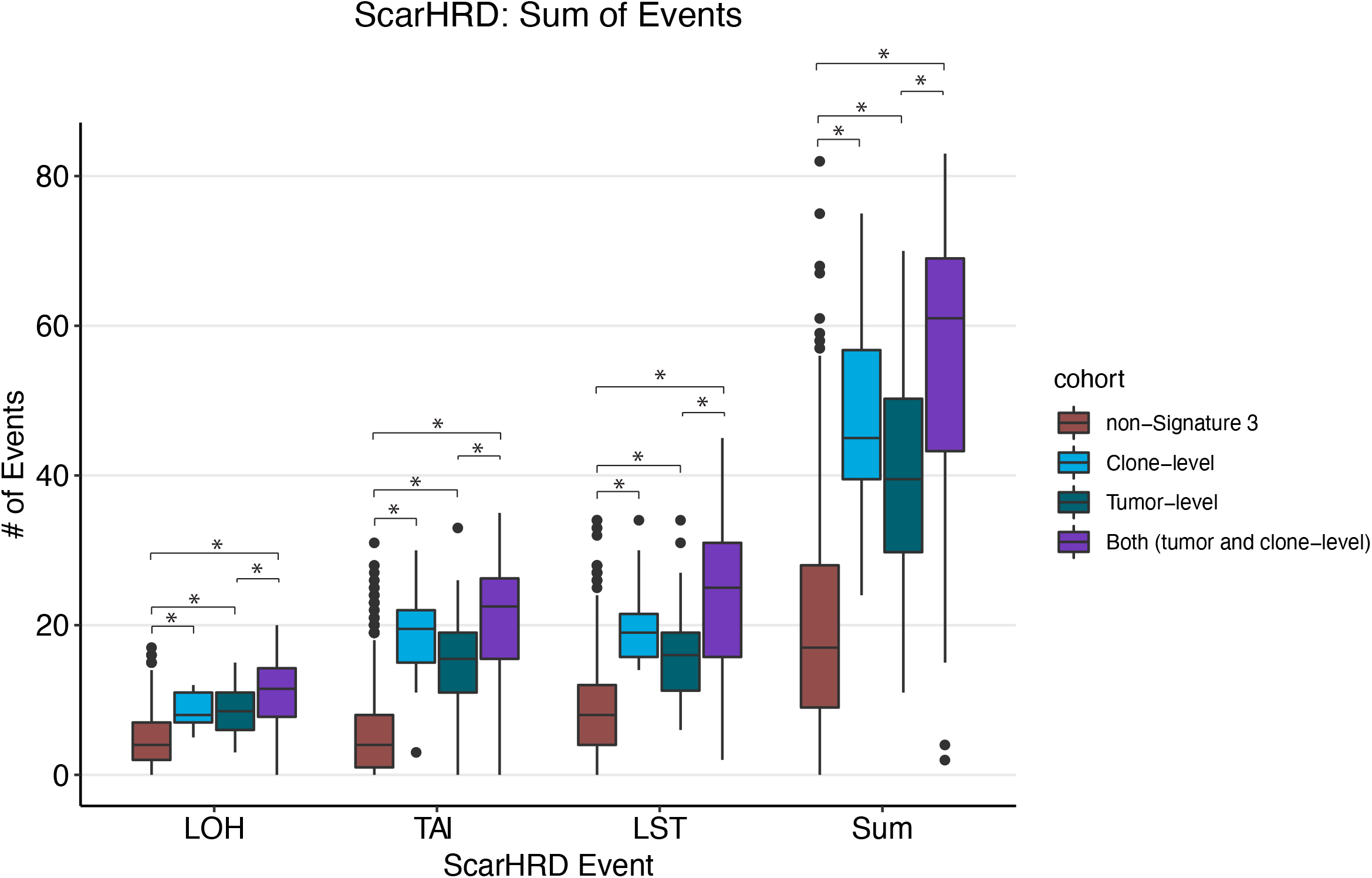
HRD-associated copy number events are associated with the clonality of mutational signature 3. The distribution of homologous recombination deficiency (HRD)-associated events between tumors where mutational signature 3 was identified at both the tumor and cell subpopulation levels, just at the tumor level, just at the cell subpopulation level, and not at all. Asterisks indicate statistical significance via Mann-Whitney U tests.

**Supplementary Figure 6:**
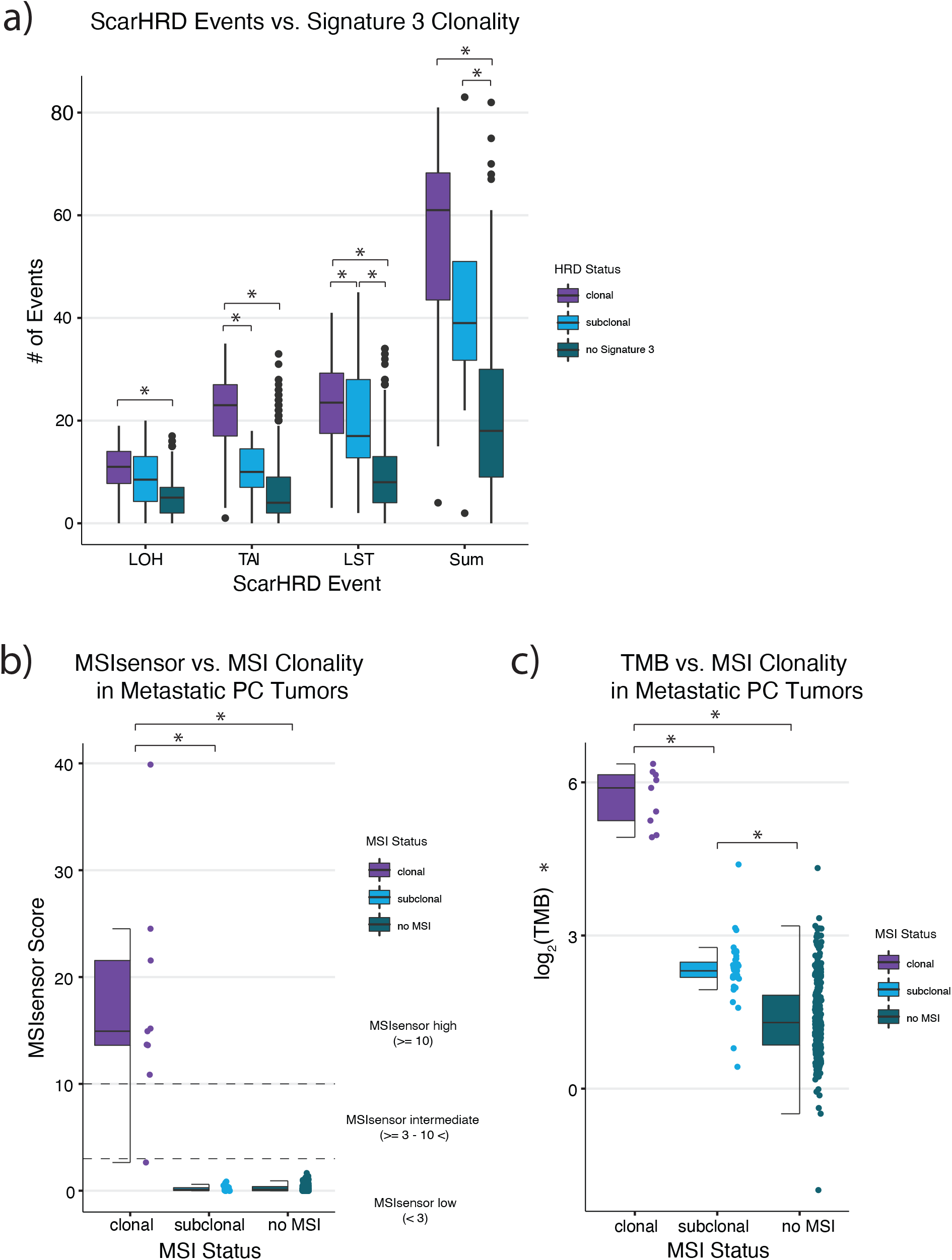
Clonality of mutational signatures and their associations with genomic events in PC tumors. **(a)** The distribution HRD-associated copy number events based on the clonality of mutational signature 3 from running SigMA at the cell subpopulation level. There is a stepwise increase in the number of HRD-associated copy number events from tumors with no signature 3, to subclonal signature 3, to clonal signature 3. **(b)** The distribution of MSIsensor scores based on the clonality of MSI-associated mutational signatures from running SigMA at the cell subpopulation level. Samples with clonal activity of MSI-associated mutational signatures had significantly higher MSIsensor scores than samples without activity of MSI-associated mutational signatures, however, this was not the case for samples with subclonal activity of MSI-associated mutational signatures. **(c)** The distribution of tumor mutational burden based on the clonality of MSI-associated mutational signatures from running SigMA at the cell subpopulation level. There is a stepwise increase in the TMB from tumors with no MSI-associated signature, to subclonal MSI-associated signature, to clonal MSI associated signature.

**Supplementary Figure 7:**
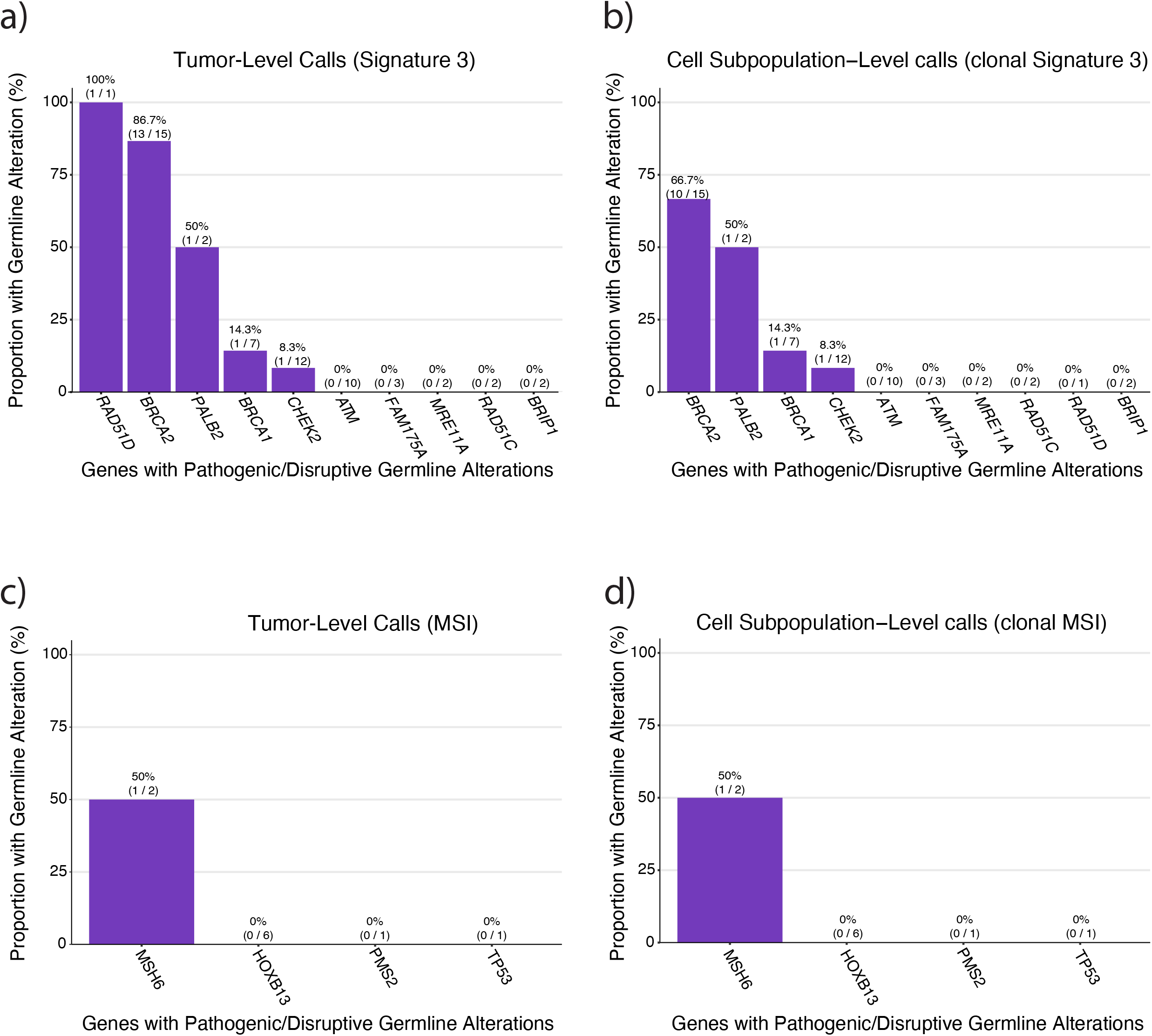
Germline and putative loss-of-function somatic alterations in DNA repair genes in samples with signature 3 and MSI. **(a)** The proportion of samples with germline alterations in genes associated with homologous recombination that exhibited mutational signature 3 when running SigMA on all mutations in the tumor and on **(b)** mutations from each cell subpopulation. Samples with germline alterations in ATM, FAM175A, MRE11A, RAD51C, and BRIP1 never exhibited mutational signature 3. **(c)** The proportion of samples with germline alterations in genes associated with mismatch repair deficiency that exhibited microsatelite instability associated mutational signatures when running SigMA on all mutations in the tumor and on **(d)** mutations from each cell subpopulation. Samples with germline alterations in HOXB13, PMS2, and TP53 never exhibited microsatelite instability associated mutational signatures. Some of the genes included in our list of homologous recombination associated and mismatch repair associated genes did not have germline alterations in our cohort, and therefore are not represented in these figures.

**Supplementary Figure 8:**
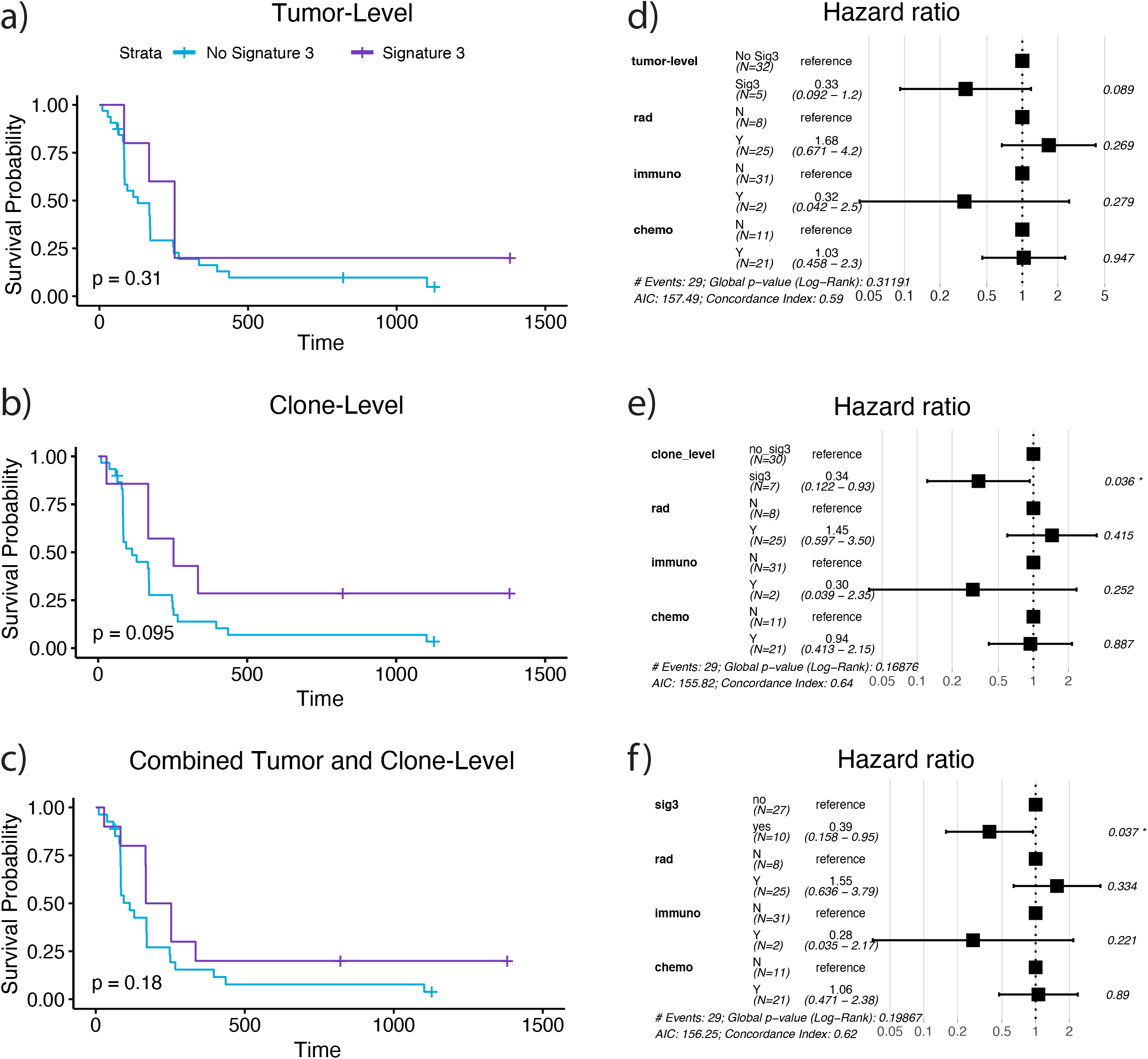
Effect of signature 3 identification framework on PARPi treated prostate cancers. **(a)** Kaplan-Meier survival curves between signature 3 and non-signature 3 tumors identified by running SigMa at the tumor-level. **(b)** Kaplan-Meier survival curves between signature 3 and non-signature 3 tumors identified by running SigMa at the cell subpopulation-level. **(c)** Kaplan-Meier survival curves between signature 3 and non-signature 3 tumors identified by combining the tumor-level and cell subpopulation-level SigMa calls. **(d)** Multivariate Cox proportional-hazards analysis for tumor-level signature 3, correcting for whether or not the patient also received radition, immune-related therapy, and chemotherapy. **(e)** Multivariate Cox proportional-hazards analysis for cell subpopulation-level signature 3, correcting for whether or not the patient also recieved radition, immune-related therapy, and chemotherapy. **(f)** Multivariate Cox proportional-hazards analysis for the combined tumor-level and cell subpopulation-level signature 3 calls, correcting for whether or not the patient also recieved radition, immune-related therapy, and chemotherapy.

**Supplementary Figure 9:**
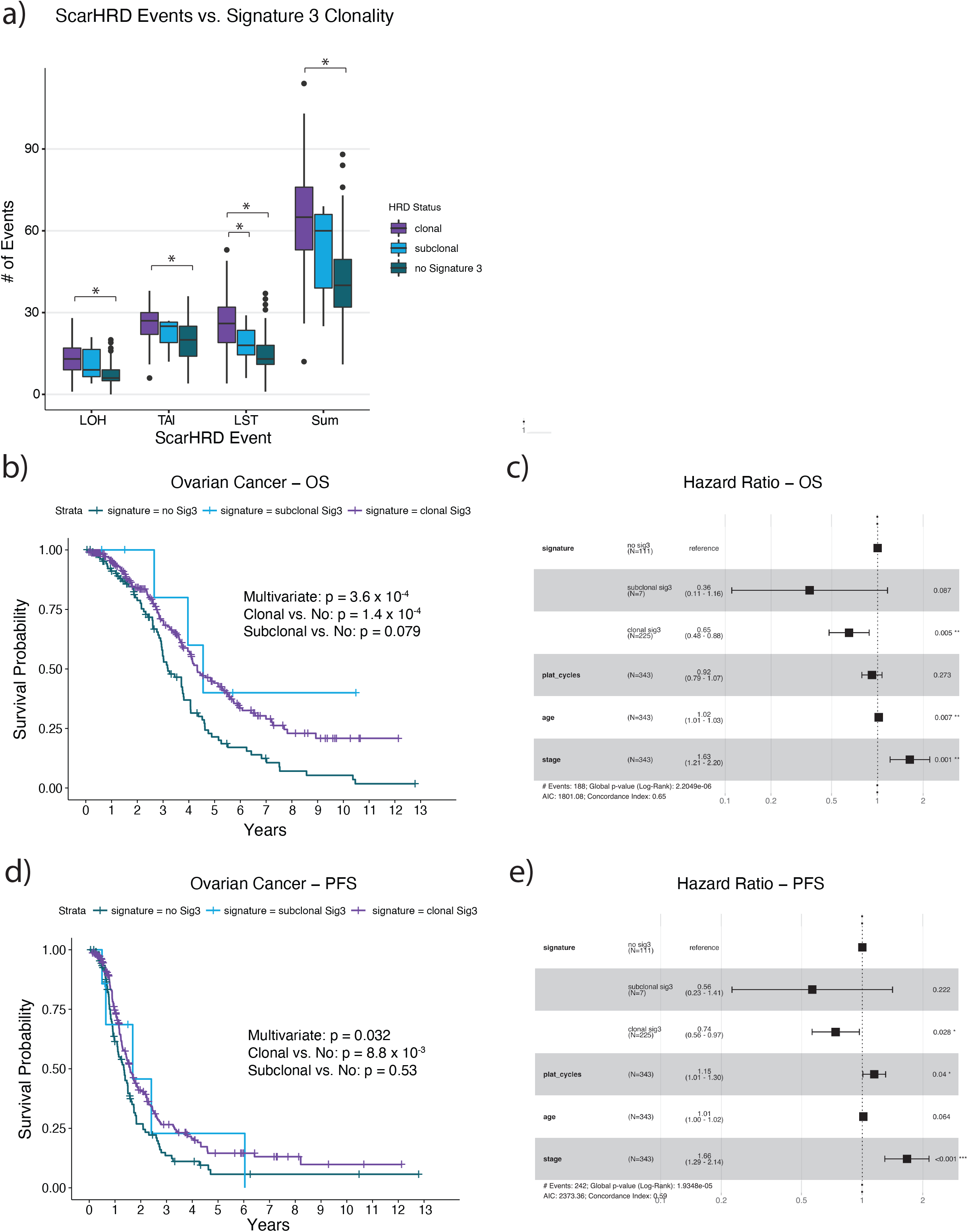
Effect of signature 3 clonality on survival in cisplatin treated ovarian cancers. **(a)** The associations between the clonality of mutational signature 3 and the number of homologous recombination deficiency (HRD)-associated copy number events in the ovarian cancer cohort were highly concordant with the observed associations in our prostate cancer cohort. That is, there is a stepwise increase in the number of HRD-associated copy number events from tumors with no signature 3 (n = 111), to subclonal signature 3 (n = 7), to clonal signature 3 (n = 225). Asterisks denote statistical significance via Mann-Whitney U tests. To determine if the clonality of mutational signature 3 effects how tumors respond to therapy, we performed Kaplan-Meier and Cox proportional hazard (PH) analysis on **(b-c)** overall survival (OS) and **(d-e)** progression free survival (PFS). Tumors with clonal activity of mutational signature 3 were associated with significantly improved **(b-c)** OS and **(d-e)** PFS. **(c)** Tumors with only subclonal activity of mutational signature were borderline significantly associated with improved OS (Cox PH, p = 0.087). **(e)** Although nonsignificant, subclonal only activity of mutational signature 3 trended in the direction of improved PFS as well (Cox PH, HR = 0.56, p = 0.22).

## References

1. Liu J, Dang H, Wang XW. The significance of intertumor and intratumor heterogeneity in liver cancer. Exp Mol Med. 2018;50:e416.

2. Roth A, Khattra J, Yap D, Wan A, Laks E, Biele J, et al. PyClone: statistical inference of clonal population structure in cancer. Nat Methods. 2014;11:396–8.

3. Andor N, Harness JV, Müller S, Mewes HW, Petritsch C. EXPANDS: expanding ploidy and allele frequency on nested subpopulations. Bioinformatics. 2014;30:50–60.

4. Gerstung M, Jolly C, Leshchiner I, Dentro SC, Gonzalez S, Rosebrock D, et al. The evolutionary history of 2,658 cancers. Nature. 2020;578:122–8.

5. Turajlic S, Sottoriva A, Graham T, Swanton C. Resolving genetic heterogeneity in cancer. Nat Rev Genet. 2019;20:404–16.

6. Siegel RL, Miller KD, Jemal A. Cancer statistics, 2020. CA Cancer J Clin. 2020;70:7–30.

7. Marcus L, Lemery SJ, Keegan P, Pazdur R. FDA Approval Summary: Pembrolizumab for the Treatment of Microsatellite Instability-High Solid Tumors. Clin Cancer Res. 2019;25:3753–8.

8. Hussain M, Mateo J, Fizazi K, Saad F, Shore N, Sandhu S, et al. Survival with Olaparib in Metastatic Castration-Resistant Prostate Cancer. N Engl J Med [Internet]. 2020; Available from: http://dx.doi.org/10.1056/NEJMoa2022485

9. Abida W, Patnaik A, Campbell D, Shapiro J, Bryce AH, McDermott R, et al. Rucaparib in Men With Metastatic Castration-Resistant Prostate Cancer Harboring a BRCA1 or BRCA2 Gene Alteration [Internet]. Journal of Clinical Oncology. 2020. page 3763–72. Available from: http://dx.doi.org/10.1200/jco.20.01035

10. Mateo J, Porta N, Bianchini D, McGovern U, Elliott T, Jones R, et al. Olaparib in patients with metastatic castration-resistant prostate cancer with DNA repair gene aberrations (TOPARP-B): a multicentre, open-label, randomised, phase 2 trial. Lancet Oncol. 2020;21:162–74.

11. Espiritu SMG, Liu LY, Rubanova Y, Bhandari V, Holgersen EM, Szyca LM, et al. The Evolutionary Landscape of Localized Prostate Cancers Drives Clinical Aggression. Cell. 2018;173:1003–1013.e15.

12. Armenia J, Wankowicz SAM, Liu D, Gao J, Kundra R, Reznik E, et al. The long tail of oncogenic drivers in prostate cancer. Nat Genet. 2018;50:645–51.

13. Gerhauser C, Favero F, Risch T, Simon R, Feuerbach L, Assenov Y, et al. Molecular Evolution of Early-Onset Prostate Cancer Identifies Molecular Risk Markers and Clinical Trajectories. Cancer Cell. 2018;34:996–1011.e8.

14. Cancer Stat Facts: Cancer Disparities [Internet]. [cited 2022 Nov 3]. Available from: https://seer.cancer.gov/statfacts/html/disparities.html

15. Rebbeck TR, Halbert CH, Sankar P. Genetics, epidemiology, and cancer disparities: is it black and white? J Clin Oncol. 2006;24:2164–9.

16. Nyame YA, Cooperberg MR, Cumberbatch MG, Eggener SE, Etzioni R, Gomez SL, et al. Deconstructing, Addressing, and Eliminating Racial and Ethnic Inequities in Prostate Cancer Care. Eur Urol [Internet]. 2022; Available from: http://dx.doi.org/10.1016/j.eururo.2022.03.007

17. Koga Y, Song H, Chalmers ZR, Newberg J, Kim E, Carrot-Zhang J, et al. Genomic Profiling of Prostate Cancers from Men with African and European Ancestry. Clin Cancer Res. 2020;26:4651–60.

18. Kamran SC, Xie J, Cheung ATM, Mavura MY, Song H, Palapattu EL, et al. Tumor Mutations Across Racial Groups in a Real-World Data Registry. JCO Precision Oncology [Internet]. Wolters Kluwer Health; 2021 [cited 2021 Nov 8]; Available from: http://dx.doi.org/10.1200/PO.21.00340

19. Huang FW, Mosquera JM, Garofalo A, Oh C, Baco M, Amin-Mansour A, et al. Exome Sequencing of African-American Prostate Cancer Reveals Loss-of-Function ERF Mutations [Internet]. Cancer Discovery. 2017. page 973–83. Available from: http://dx.doi.org/10.1158/2159-8290.cd-16-0960

20. Petrovics G, Li H, Stümpel T, Tan S-H, Young D, Katta S, et al. A novel genomic alteration of LSAMP associates with aggressive prostate cancer in African American men. EBioMedicine. 2015;2:1957–64.

21. Stopsack KH, Nandakumar S, Arora K, Nguyen B, Vasselman SE, Nweji B, et al. Differences in Prostate Cancer Genomes by Self-reported Race: Contributions of Genetic Ancestry, Modifiable Cancer Risk Factors, and Clinical Factors. Clin Cancer Res. American Association for Cancer Research; 2022;28:318–26.

22. Di Rienzo Agustin Fuentes Stephanie M. Fullerton Nanibaa’ A. Garrison Nayanika Ghosh Evelynn M. Hammonds David S. Jones Eimear E. Kenny Peter Kraft Sandra S.-J. Lee Madelyn Mauro John Novembre Aaron Panofsky Mashaal Sohail Benjamin M. Neale Danielle S. Allen ACFLSJMPSABDA. Getting genetic ancestry right for science and society. Science [Internet]. 2022;376. Available from: http://dx.doi.org/10.1126/science.abm7530

23. Wedge DC, Gundem G, Mitchell T, Woodcock DJ, Martincorena I, Ghori M, et al. Sequencing of prostate cancers identifies new cancer genes, routes of progression and drug targets. Nat Genet. 2018;50:682–92.

24. Crumbaker M, Khoja L, Joshua AM. AR Signaling and the PI3K Pathway in Prostate Cancer. Cancers [Internet]. 2017;9. Available from: http://dx.doi.org/10.3390/cancers9040034

25. Bitting RL, Armstrong AJ. Targeting the PI3K/Akt/mTOR pathway in castration-resistant prostate cancer. Endocr Relat Cancer. 2013;20:R83–99.

26. Alexandrov LB, Kim J, Haradhvala NJ, Huang MN, Tian Ng AW, Wu Y, et al. The repertoire of mutational signatures in human cancer. Nature. 2020;578:94–101.

27. Alexandrov LB, Nik-Zainal S, Wedge DC, Aparicio SAJR, Behjati S, Biankin AV, et al. Signatures of mutational processes in human cancer. Nature. 2013;500:415–21.

28. Gulhan DC, Lee JJ-K, Melloni GEM, Cortés-Ciriano I, Park PJ. Detecting the mutational signature of homologous recombination deficiency in clinical samples. Nat Genet. 2019;51:912–9.

29. Mateo J, Carreira S, Sandhu S, Miranda S, Mossop H, Perez-Lopez R, et al. DNA-Repair Defects and Olaparib in Metastatic Prostate Cancer. N Engl J Med. 2015;373:1697–708.

30. Pritchard CC, Mateo J, Walsh MF, De Sarkar N, Abida W, Beltran H, et al. Inherited DNA-Repair Gene Mutations in Men with Metastatic Prostate Cancer. N Engl J Med. 2016;375:443–53.

31. de Bono J, Kang J, Hussain M. Olaparib for Metastatic Castration-Resistant Prostate Cancer. Reply. N. Engl. J. Med. 2020. page 891.

32. Sztupinszki Z, Diossy M, Krzystanek M, Reiniger L, Csabai I, Favero F, et al. Migrating the SNP array-based homologous recombination deficiency measures to next generation sequencing data of breast cancer. NPJ Breast Cancer. 2018;4:16.

33. Niu B, Ye K, Zhang Q, Lu C, Xie M, McLellan MD, et al. MSIsensor: microsatellite instability detection using paired tumor-normal sequence data. Bioinformatics. 2014;30:1015–6.

34. Akdemir KC, Le VT, Kim JM, Killcoyne S, King DA, Lin Y-P, et al. Somatic mutation distributions in cancer genomes vary with three-dimensional chromatin structure. Nat Genet. 2020;52:1178–88.

35. Ma J, Setton J, Lee NY, Riaz N, Powell SN. The therapeutic significance of mutational signatures from DNA repair deficiency in cancer. Nat Commun. 2018;9:3292.

36. Sztupinszki Z, Diossy M, Krzystanek M, Borcsok J, Pomerantz MM, Tisza V, et al. Detection of Molecular Signatures of Homologous Recombination Deficiency in Prostate Cancer with or without BRCA1/2 Mutations. Clin Cancer Res. American Association for Cancer Research; 2020;26:2673–80.

37. Abida W, Cyrta J, Heller G, Prandi D, Armenia J, Coleman I, et al. Genomic correlates of clinical outcome in advanced prostate cancer. Proc Natl Acad Sci U S A. 2019;116:11428– 36.

38. Network TCGAR, The Cancer Genome Atlas Research Network. Integrated genomic analyses of ovarian carcinoma [Internet]. Nature. 2011. page 609–15. Available from: http://dx.doi.org/10.1038/nature10166

39. Ellrott K, Bailey MH, Saksena G, Covington KR, Kandoth C, Stewart C, et al. Scalable Open Science Approach for Mutation Calling of Tumor Exomes Using Multiple Genomic Pipelines. Cell Syst. 2018;6:271–281.e7.

40. Rosenthal R, McGranahan N, Herrero J, Taylor BS, Swanton C. DeconstructSigs: delineating mutational processes in single tumors distinguishes DNA repair deficiencies and patterns of carcinoma evolution. Genome Biol. 2016;17:31.

41. Blokzijl F, Janssen R, van Boxtel R, Cuppen E. MutationalPatterns: comprehensive genome-wide analysis of mutational processes. Genome Med. 2018;10:33.

42. Gopal P, Sarihan EI, Chie EK, Kuzmishin G, Doken S, Pennell NA, et al. Clonal selection confers distinct evolutionary trajectories in BRAF-driven cancers. Nat Commun. 2019;10:5143.

43. Notta F, Chan-Seng-Yue M, Lemire M, Li Y, Wilson GW, Connor AA, et al. A renewed model of pancreatic cancer evolution based on genomic rearrangement patterns. Nature. 2016;538:378–82.

44. Berchuck_lipids_cistrome.pdf [Internet]. Available from: http://dx.doi.org/10.1158/0008-5472.CAN-21-3552/3164400/can-21-3552.pdf

45. Turajlic S, Xu H, Litchfield K, Rowan A, Horswell S, Chambers T, et al. Deterministic Evolutionary Trajectories Influence Primary Tumor Growth: TRACERx Renal. Cell. 2018;173:595–610.e11.

46. Powell IJ, Bock CH, Ruterbusch JJ, Sakr W. Evidence supports a faster growth rate and/or earlier transformation to clinically significant prostate cancer in black than in white American men, and influences racial progression and mortality disparity. J Urol. 2010;183:1792–6.

47. Dess RT, Hartman HE, Mahal BA, Soni PD, Jackson WC, Cooperberg MR, et al. Association of Black Race With Prostate Cancer-Specific and Other-Cause Mortality. JAMA Oncol. 2019;5:975–83.

48. George DJ, Ramaswamy K, Huang A, Russell D, Mardekian J, Schultz NM, et al. Survival by race in men with chemotherapy-naive enzalutamide- or abiraterone-treated metastatic castration-resistant prostate cancer. Prostate Cancer Prostatic Dis [Internet]. 2021; Available from: http://dx.doi.org/10.1038/s41391-021-00463-9

49. Spratt DE, Chen Y-W, Mahal BA, Osborne JR, Zhao SG, Morgan TM, et al. Individual Patient Data Analysis of Randomized Clinical Trials: Impact of Black Race on Castration-resistant Prostate Cancer Outcomes. Eur Urol Focus. 2016;2:532–9.

50. Haraldsdottir S, Hampel H, Tomsic J, Frankel WL, Pearlman R, de la Chapelle A, et al. Colon and endometrial cancers with mismatch repair deficiency can arise from somatic, rather than germline, mutations. Gastroenterology. 2014;147:1308–1316.e1.

51. Abida W, Cheng ML, Armenia J, Middha S, Autio KA, Vargas HA, et al. Analysis of the Prevalence of Microsatellite Instability in Prostate Cancer and Response to Immune Checkpoint Blockade. JAMA Oncol. 2019;5:471–8.

52. de Bono J, Mateo J, Fizazi K, Saad F, Shore N, Sandhu S, et al. Olaparib for Metastatic Castration-Resistant Prostate Cancer. N Engl J Med. 2020;382:2091–102.

53. Polak P, Kim J, Braunstein LZ, Karlic R, Haradhavala NJ, Tiao G, et al. A mutational signature reveals alterations underlying deficient homologous recombination repair in breast cancer. Nat Genet. 2017;49:1476–86.

54. Turajlic S, Xu H, Litchfield K, Rowan A, Chambers T, Lopez JI, et al. Tracking Cancer Evolution Reveals Constrained Routes to Metastases: TRACERx Renal. Cell. 2018;173:581–594.e12.

55. Jamal-Hanjani M, Wilson GA, McGranahan N, Birkbak NJ, Watkins TBK, Veeriah S, et al. Tracking the Evolution of Non-Small-Cell Lung Cancer. N Engl J Med. 2017;376:2109–21.

56. Kim C, Gao R, Sei E, Brandt R, Hartman J, Hatschek T, et al. Chemoresistance Evolution in Triple-Negative Breast Cancer Delineated by Single-Cell Sequencing. Cell. 2018;173:879–893.e13.

57. Casasent AK, Schalck A, Gao R, Sei E, Long A, Pangburn W, et al. Multiclonal Invasion in Breast Tumors Identified by Topographic Single Cell Sequencing. Cell. 2018;172:205–217.e12.

58. Zhang K. Stratifying tissue heterogeneity with scalable single-cell assays. Nat. Methods. 2017. page 238–9.

59. Zheng GXY, Lau BT, Schnall-Levin M, Jarosz M, Bell JM, Hindson CM, et al. Haplotyping germline and cancer genomes with high-throughput linked-read sequencing. Nat Biotechnol. 2016;34:303–11.

60. Carter SL, Cibulskis K, Helman E, McKenna A, Shen H, Zack T, et al. Absolute quantification of somatic DNA alterations in human cancer. Nat Biotechnol. 2012;30:413– 21.

61. Shen R, Seshan VE. FACETS: allele-specific copy number and clonal heterogeneity analysis tool for high-throughput DNA sequencing. Nucleic Acids Res. 2016;44:e131.

62. Robinson D, Van Allen EM, Wu Y-M, Schultz N, Lonigro RJ, Mosquera J-M, et al. Integrative clinical genomics of advanced prostate cancer. Cell. 2015;161:1215–28.

63. Cancer Genome Atlas Research Network. The Molecular Taxonomy of Primary Prostate Cancer. Cell. 2015;163:1011–25.

64. Beltran H, Prandi D, Mosquera JM, Benelli M, Puca L, Cyrta J, et al. Divergent clonal evolution of castration-resistant neuroendocrine prostate cancer. Nat Med. 2016;22:298– 305.

65. Kumar A, Coleman I, Morrissey C, Zhang X, True LD, Gulati R, et al. Substantial interindividual and limited intraindividual genomic diversity among tumors from men with metastatic prostate cancer. Nat Med. 2016;22:369–78.

66. Barbieri CE, Baca SC, Lawrence MS, Demichelis F, Blattner M, Theurillat J-P, et al. Exome sequencing identifies recurrent SPOP, FOXA1 and MED12 mutations in prostate cancer. Nat Genet. 2012;44:685–9.

67. Baca SC, Prandi D, Lawrence MS, Mosquera JM, Romanel A, Drier Y, et al. Punctuated evolution of prostate cancer genomes. Cell. 2013;153:666–77.

68. Huang FW, Mosquera JM, Garofalo A, Oh C, Baco M, Amin-Mansour A, et al. Exome Sequencing of African-American Prostate Cancer Reveals Loss-of-Function Mutations. Cancer Discov. 2017;7:973–83.

69. McGranahan N, Favero F, de Bruin EC, Birkbak NJ, Szallasi Z, Swanton C. Clonal status of actionable driver events and the timing of mutational processes in cancer evolution. Sci Transl Med. 2015;7:283ra54.

70. Sun Y, Meng R, Cheng Z-Y, Fan C, Wei X-M, Yang Y, et al. Characterization of genomic clones using circulating tumor DNA in patients with hepatocarcinoma [Internet]. Translational Cancer Research. 2018. page 321–9. Available from: http://dx.doi.org/10.21037/tcr.2018.03.17

71. Lawrence MS, Stojanov P, Polak P, Kryukov GV, Cibulskis K, Sivachenko A, et al. Mutational heterogeneity in cancer and the search for new cancer-associated genes. Nature. 2013;499:214–8.

72. Abkevich V, Timms KM, Hennessy BT, Potter J, Carey MS, Meyer LA, et al. Patterns of genomic loss of heterozygosity predict homologous recombination repair defects in epithelial ovarian cancer. Br J Cancer. 2012;107:1776–82.

73. Birkbak NJ, Wang ZC, Kim J-Y, Eklund AC, Li Q, Tian R, et al. Telomeric allelic imbalance indicates defective DNA repair and sensitivity to DNA-damaging agents. Cancer Discov. 2012;2:366–75.

74. Popova T, Manié E, Rieunier G, Caux-Moncoutier V, Tirapo C, Dubois T, et al. Ploidy and large-scale genomic instability consistently identify basal-like breast carcinomas with BRCA1/2 inactivation. Cancer Res. 2012;72:5454–62.

75. Kamburov A, Wierling C, Lehrach H, Herwig R. ConsensusPathDB—a database for integrating human functional interaction networks [Internet]. Nucleic Acids Research. 2009. page D623–8. Available from: http://dx.doi.org/10.1093/nar/gkn698

76. Kamburov A, Pentchev K, Galicka H, Wierling C, Lehrach H, Herwig R. ConsensusPathDB: toward a more complete picture of cell biology [Internet]. Nucleic Acids Research. 2011. page D712–7. Available from: http://dx.doi.org/10.1093/nar/gkq1156

77. Poplin R, Chang P-C, Alexander D, Schwartz S, Colthurst T, Ku A, et al. A universal SNP and small-indel variant caller using deep neural networks [Internet]. Nature Biotechnology. 2018. page 983–7. Available from: http://dx.doi.org/10.1038/nbt.4235

78. AlDubayan SH, Conway JR, Camp SY, Witkowski L, Kofman E, Reardon B, et al. Detection of Pathogenic Variants With Germline Genetic Testing Using Deep Learning vs Standard Methods in Patients With Prostate Cancer and Melanoma. JAMA. 2020;324:1957–69.

79. DePristo MA, Banks E, Poplin R, Garimella KV, Maguire JR, Hartl C, et al. A framework for variation discovery and genotyping using next-generation DNA sequencing data. Nat Genet. 2011;43:491–8.

80. Tan A, Abecasis GR, Kang HM. Unified representation of genetic variants. Bioinformatics. 2015;31:2202–4.

81. McLaren W, Gil L, Hunt SE, Riat HS, Ritchie GRS, Thormann A, et al. The Ensembl Variant Effect Predictor. Genome Biol. 2016;17:122.

82. Richards S, Aziz N, Bale S, Bick D, Das S, Gastier-Foster J, et al. Standards and guidelines for the interpretation of sequence variants: a joint consensus recommendation of the American College of Medical Genetics and Genomics and the Association for Molecular Pathology. Genet Med. 2015;17:405–24.

83. Consortium T 1000 GP, The 1000 Genomes Project Consortium. A global reference for human genetic variation [Internet]. Nature. 2015. page 68–74. Available from: http://dx.doi.org/10.1038/nature15393

84. Clarke L, Fairley S, Zheng-Bradley X, Streeter I, Perry E, Lowy E, et al. The international Genome sample resource (IGSR): A worldwide collection of genome variation incorporating the 1000 Genomes Project data. Nucleic Acids Res. 2017;45:D854–9.

85. Chang CC, Chow CC, Tellier LC, Vattikuti S, Purcell SM, Lee JJ. Second-generation PLINK: rising to the challenge of larger and richer datasets [Internet]. GigaScience. 2015. Available from: http://dx.doi.org/10.1186/s13742-015-0047-8

86. Jørsboe E, Hanghøj K, Albrechtsen A. fastNGSadmix: admixture proportions and principal component analysis of a single NGS sample. Bioinformatics. 2017;33:3148–50.

